# Two faces of perceptual awareness during the attentional blink: Gradual and discrete

**DOI:** 10.1101/769083

**Authors:** Aytaç Karabay, Sophia A. Wilhelm, Joost de Jong, Jing Wang, Sander Martens, Elkan G. Akyürek

**Affiliations:** Department of Psychology, Experimental Psychology, University of Groningen, The Netherlands; Brain Function and Psychological Science Research Center, Shenzhen University, China; Cognitive Neuroscience Center, University of Groningen, The Netherlands; Department of Biomedical Sciences of Cells & Systems, University Medical Center Groningen, The Netherlands

**Keywords:** attentional blink, perceptual awareness, working memory, continuous report, precision, guess rate

## Abstract

In a series of experiments, the nature of perceptual awareness during the attentional blink was investigated. Previous work has considered the attentional blink as a discrete, all-or-none phenomenon, indicative of general access to conscious awareness. Using continuous report measures in combination with mixture modeling, the outcomes showed that perceptual awareness during the attentional blink can be a gradual phenomenon. Awareness was not exclusively discrete, but also exhibited a gradual characteristic whenever the spatial extent of attention induced by the first target spanned more than a single location. Under these circumstances, mental representations of blinked targets were impoverished, but did approach the actual identities of the targets. Conversely, when the focus of attention covered only a single location, there was no evidence for any partial knowledge of blinked targets. These two different faces of awareness during the attentional blink challenge current theories of both awareness and temporal attention, which cannot explain the existence of gradual awareness of targets during the attentional blink. To account for the current outcomes, an adaptive gating model is proposed that casts awareness on a continuum between gradual and discrete, rather than as being of either single kind.

---

Attention is a critical cognitive function. Through the attentional selection of objects and events of interest, at the expense of other items that appear less interesting, cognition can be devoted to information that is likely to be relevant. Attention strongly drives human behavior: We act on what we select and process cognitively, not on things that are ignored. Although certain basic properties of an object or event might be registered by the brain independent of attention (e.g., a global motion path), attention generally limits entry to higher-level processing, as is needed to actually identify and name an object. On top of that, selective attention is also seen as the gatekeeper of conscious awareness (Dehaene & Changeux, 2011; Mashour et al., 2020), which is similarly concerned with only one or a few items at any one time (e.g., Posner, 1994).

The selective nature of attention is both its strength and its weakness. Its selectivity assures that not everything that is sensed is subsequently processed in-depth, allowing major savings in cognitive expenditure. However, attentional selectivity also implies a limited ability that can fall short when there are more than a few items of interest present. It is therefore not surprising that failures of attention are thought to contribute to human error (Endsley, 1995). A compelling demonstration of the very limited nature of attention is the attentional blink (AB) phenomenon. The AB is the difficulty in selecting a task-relevant target stimulus if it occurs within approximately half a second after another target (Broadbent & Broadbent, 1987; Raymond et al., 1992). Without selecting that prior target, or with sufficient time in-between, no difficulty is experienced. Thus, regardless of the specific theoretical interpretation (for an overview, see Dux & Marois, 2009; Martens & Wyble, 2010), it is evident that the attentional processing of just a single preceding target already strongly impairs the processing of the following one, and does so for a considerable length of time.

The AB phenomenon has proven to be extremely robust; it occurs in the vast majority of people, and is observed in diverse tasks (Martens & Wyble, 2010). The most common of these tasks is the rapid serial visual presentation (RSVP) paradigm, in which stimuli are presented at a rate of approximately 10 per second. Within a sequential stream of distractor items, two targets are inserted at variable delays from each other (called “lag”). Observers are asked to identify both targets, and the AB is reflected in their reduced ability to successfully report the second target (T2) when the lag from the first target (T1) is short. Although classic RSVP tasks present all stimuli at a single, central location, the AB has also been observed in tasks that test performance across the visual field. The oldest and most well-known task to do so is the dwell time (DT) paradigm, pioneered by Duncan, Ward, and Shapiro (1994). In this task, one target appeared at either the left or right of a central fixation dot, while the other target appeared either above or below fixation. Instead of a distractor stream, the targets were each followed by a single pattern mask. Despite these procedural differences, as in RSVP, with short delays between targets, the identification of the trailing target was impaired, evidence of the AB.

The universal nature of the AB suggests that it is the consequence of a general property of cognition. One particularly influential conceptualization of that idea is that the AB quite directly reflects what is being perceived consciously, and what is not. In the Global Neuronal Workspace model of conscious access proposed by Dehaene, Sergent, and Changeux (2003; cf. Baars, 1989), conscious awareness is associated with items that mobilize a global, long-distance network in the brain, through which their representations become available to various cognitive processes (e.g., verbal report, memory). Items that have not (yet) entered conscious awareness can at most receive local processing, for instance in occipital regions, where basic visual features may be extracted, but not much more. As an item enters the global workspace, initially driven by bottom-up neural activity that eventually ‘ignites’ the global workspace, the neurons that carry the item’s signal inhibit other neurons that might propagate other items, thereby preventing them from entering awareness.

In terms of the AB, the Global Neuronal Workspace model predicts the following. T1 is logically the first to be selected, to enter the global workspace, and to gain conscious access. It then inhibits T2 from subsequently entering the workspace. Therefore, T2 is relegated to lower-level areas (e.g., primary sensory cortices), and cannot reach consciousness. Importantly, this implies an all-or-none situation; if T2 is blinked and denied access to the global workspace, its low-level activity traces cannot be consolidated (episodically) and will inevitably perish, so that the identity of T2 is utterly lost. Sergent and Dehaene (2004) verified this prediction by asking participants to report on the visibility of targets using a continuous response scale. Participants indeed tended towards either indicating no visibility at all for blinked T2s, or high visibility for T2s that were not blinked, with virtually no evidence for cases of partial awareness.

In addition to subjective judgments about perceptual clarity that may be considered unreliable and/or prone to bias (e.g., Hannula et al., 2005), there is also electrophysiological evidence in support of an all-or-none characteristic of conscious awareness during the AB. Event-related potential (ERP) studies of the AB demonstrated that even though early perceptual ERP components (the P1 and N1) survive the AB, the N2pc and the P3 are strongly suppressed (Dell’Acqua et al., 2006; Kranczioch et al., 2003; Luck et al., 1996; Vogel et al., 1998; Sergent et al., 2005). The P3, in particular, has been linked to memory consolidation and to the generation of response decisions (Donchin & Coles, 1988; Kok, 2001; Polich, 2007; Verleger et al., 2005), exactly the kind of processes that would be expected to suffer from the AB according to the Global Neuronal Workspace theory (Dehaene, Sergent, and Changeux, 2003; Sergent & Dehaene, 2004).

One might argue at this point that the typical tasks used to measure the AB present observers with categorical targets (e.g., letters), for which it is inherently difficult to measure any gradual, partial representation. Either the representation is strong enough for proper identification, or it is not. In other words, categorical identification implies having to use a certain threshold function, which naturally produces a binary outcome, and which may even influence subjective visibility reports. To more objectively assess whether there is any perceptual evidence surviving the AB, such a categorical threshold should be avoided. It is possible to do so by having observers respond to continuous target properties, such as its color [footnote 1]. The deviation in the responses as compared to the target identities can then be assessed and classified in a mixture model of the responses in feature space. In such a model, response errors fall into two distributions, a uniform distribution indicating the guess rate, and a von Mises distribution (for circular response scales) on top of the uniform distribution indicating precision (Figure 1a). Starting with the first, if the target does not reach perceptual awareness, responses should not be correlated with the target value. Hence, these random responses should form a uniform, flat distribution, which reflects the guess rate. If the target reaches perceptual awareness, to a degree, there should be a distribution of responses clustering around the target value, the standard deviation of which then shows the precision of the response (Zhang & Luck, 2008; Bays et al., 2009).

**Figure 1.**
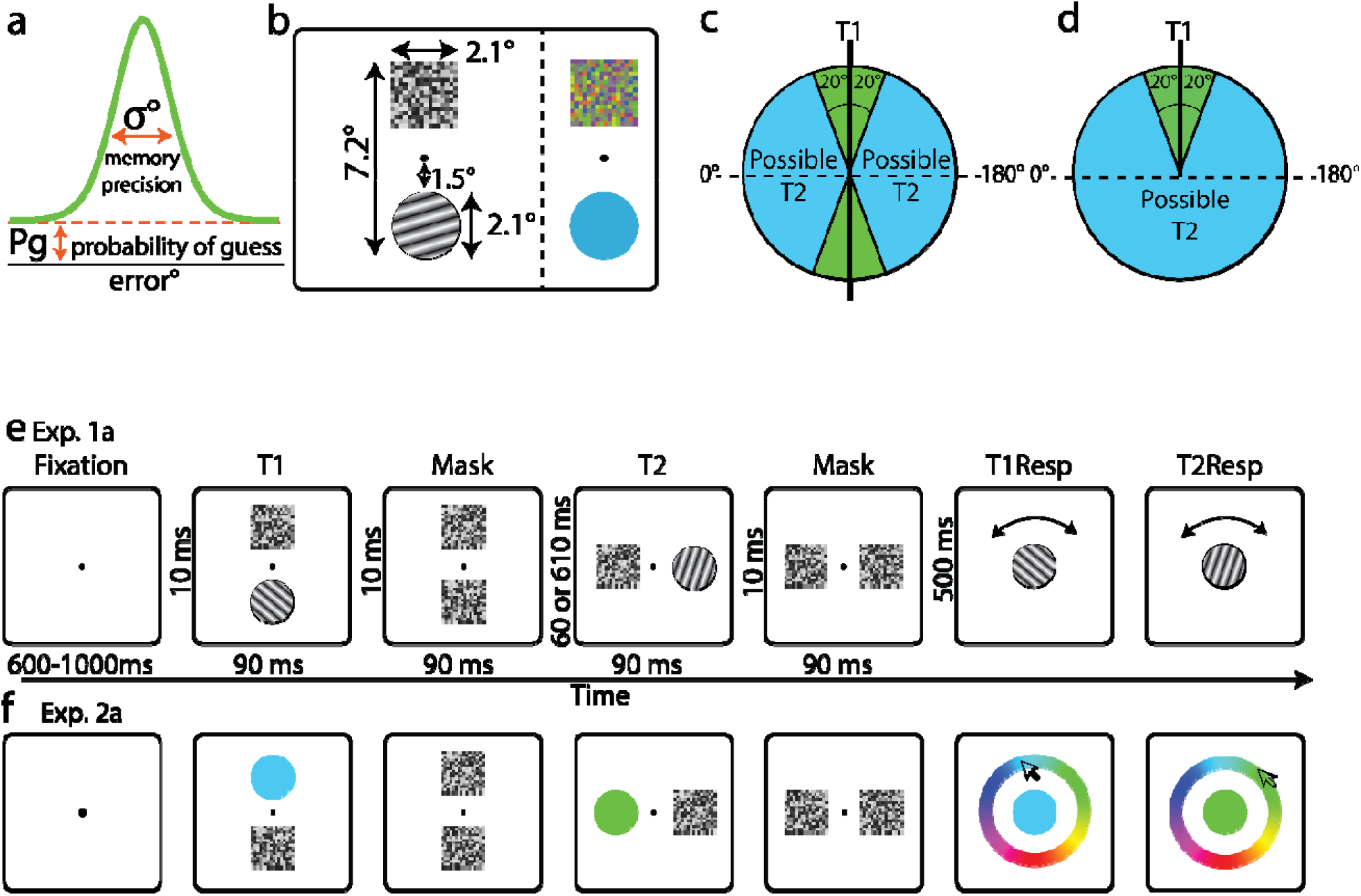
The mixture model, stimulus properties, and experimental tasks used in Experiment 1 and 2. **a** An illustration of mixture models. Pg is the proportion of the uniform distribution indicating the probability of guesses for a given condition, and σ is the standard deviation of the von Mises distribution reported in degrees (°), which indicates the precision of memory. **b** Stimulus size and appearance. Degrees (°) are given in visual angle. **c** Possible T2 orientation relative to T1 orientation, shown for a 90° T1. **d** Possible T2 color from the wheel relative to T1 color shown at 90°. **e** Trial schematic of Experiment 1. T1 means Target 1 and T2 means Target 2. T1Resp indicates T1 response prompt and T2Resp indicates T2 response prompt. f Trial schematic of Experiment 2A. Stimulus timing was identical to Experiment 1.

Asplund, Fougnie, Zughni, Martin, and Marois (2014) implemented such continuous reports in a classic RSVP paradigm, by including a T2 whose color the observers had to indicate by using a color wheel. They observed that the AB was expressed in an exclusive effect on guess rate, not on precision. The authors concluded that when the AB occurred, no representation of T2 remained at all, and any response was simply a guess. Conversely, when there was no AB, T2 was perceived as accurately as T1. There was no evidence for cases in which an impoverished representation of T2, approximating the actual T2 color, might have been salvaged from the AB. These outcomes thus provide strong evidence for an all-or-none property of the AB.

This is not to say that no target-related information can escape the AB, however. Certain stimulus characteristics do remain accessible when the AB occurs. Notably, semantic information seems to be spared, as the N400 component of the ERP (Kutas & Hillyard, 1980; Kutas & Federmeier, 2011), which reflects such relatively advanced stimulus processing, is not reduced by the AB (Luck et al., 1996; Sergent et al., 2005; though see also Giesbrecht et al., 2007). Response-related information may also survive and prime subsequent behavior (Akyürek & Hommel, 2007). Although such partial information surviving the AB may seem problematic for all-or-none models, it can be accounted for by ongoing, mostly low-level processing that might proceed without engaging attention or conscious awareness. Similarly, if it is assumed that awareness can arise at the feature level, rather than the object level, all-or-none models can also explain the finding that one feature of a target can be blinked, while another is not (Elliott, Baird, & Giesbrecht, 2016).

How such an all-or-none account might work in practice is illustrated by one of the most advanced and versatile models of the AB, the eSTST model by Wyble, Bowman, and Nieuwenstein (2009). eSTST was originally based on the two-stage model (Chun & Potter, 1995) and the idea of types and tokens (Kanwisher, 1987). In the model, types are generic stimulus representations, while tokens are their episodic (integrated) registrations. In eSTST, attention is triggered by bottom-up activation from the incoming stimuli, starting with T1. When attention is sufficiently elicited, T1 can enter the binding pool, in which types are bound to tokens, and which is situated in working memory. The ongoing binding/consolidation process inhibits attention, which keeps T2 from entering working memory. In the model, this inhibition serves the specific purpose of keeping targets apart episodically. Thus, when the AB occurs, T1 is episodically represented, consolidated, and therefore available for report, but T2 is not. However, a type-based representation of T2 may still be available, at least for a few hundred milliseconds, which might incorporate featural and even semantic as well as response-related properties.

It would thus seem that the case for discrete, all-or-none models of conscious access in the AB is very strong. Nevertheless, not all theories of conscious awareness assume it is discrete (e.g., Overgaard et al., 2006; Ramsøy & Overgaard, 2004; Sandberg et al., 2010). Furthermore, all of the evidence in favor of discrete awareness that was collected to date has come from RSVP studies, in which everything happens at a single location. It is well known that space is special in visual perception, not only because the primary visual cortex is organized accordingly (Hubel & Wiesel, 1974), but also because it seems essential for most episodic object representations to possess a “where” property. However, spatial variability, or the lack thereof, has never been considered much of a critical issue in AB research (e.g., Lunau & Olivers, 2010), and it has often been tacitly assumed that the AB is a central phenomenon that is similarly manifest in RSVP as in other tasks such as the DT paradigm (e.g., Martens & Wyble, 2010).

Nevertheless, this assumption may be questioned, not only for a lack of empirical verification, but also because of previously observed discrepancies between tasks. Consider, for instance, the case of Lag 1 sparing; an AB-related phenomenon in which T2 performance is atypically high when the targets follow each other directly, without an intervening distractor (Visser et al., 1999a). Sparing is related to the temporal integration of the targets into a unified representation (Akyürek et al., 2012; Hommel & Akyürek, 2005), which may, in turn, underlie an attentional episode. Critically, sparing is abolished when there is a spatial switch between targets (Breitmeyer et al., 1999; Visser et al., 1999a, 1999b), which suggests that certain, possibly attentional, processes might have changed. Another line of evidence that space is special in the AB comes from studies that have explicitly looked at the AB across different locations (Kristjansson & Nakayama, 2002; Wyble & Swan, 2015). In these studies, the AB was not abolished across space, but there was a spatial distribution to the effect (with the strongest AB at the same target location), suggesting that there is some spatial specificity to the AB.

In view of these considerations on the role of space in the AB, it is conceivable that even though there may be an attentional limitation across time as well as space, it could be of a different nature. Elaborating on eSTST theory (Wyble et al., 2009), it may be hypothesized that episodic representations are spatially specific. If this is true, then the ‘gate-closing’ inhibitory mechanisms that serve to maintain episodic distinctiveness, may not apply when objects are spatially separated. Consequently, the all-or-none characteristic of conscious awareness in the AB may disappear.

The current study was designed to test the prediction that gradual awareness may occur during the AB by applying a continuous report measure in both DT and RSVP paradigms. To preempt the results, the collective set of experiments showed substantial evidence not only for discrete, but also for gradual awareness. Whenever attention could be focused on a single spatial location to detect T1, subsequent T2 awareness appeared to be all-or-none. However, when the focus of attention was spread more broadly in anticipation of T1, then T2 awareness had a gradual characteristic.

## Experiment 1A

As discussed, to assess the nature of perceptual awareness in the DT paradigm, a continuous target response measure is needed. To this end, line orientation gratings were introduced as targets, and participants were asked to reproduce target orientations at the end of the trial by adjusting the orientation of probe gratings. The response error distributions for different stimulus onset asynchronies (SOAs) could thus be modeled, and estimates of precision and guess rate were generated.

### Method

#### Participants

The sample size was estimated with G-power (Faul et al., 2007), using α = .05, and β = .2, as is commonly done (Lakens, 2013). A priori power computations suggested that 24 participants would be needed to detect medium to large effect sizes (Cohen’s d_z_ = .6), so this number was the goal for each experiment. Since reliable model parameter estimates cannot be obtained with fewer than 50 data points (Bays et al., 2009), participants with less than that in any of the experimental conditions were excluded from the analysis. Participants who clearly failed to perform the task correctly, that is, who performed at or near chance level on either T1 or T2|T1 (average error > 40°), were also excluded from further analyses. In total, 30 students participated in Experiment 1, 7 of which were excluded on the basis of these criteria. In the final sample, there were thus 23 students (16 females, 7 males) at the University of Groningen, who participated in in exchange for course credits (mean age = 20.2, range = 18-24). All participants reported (corrected-to-) normal vision. The study was approved by the ethical committee of the Psychology Department of the University of Groningen (research code 18254-SP). All participants signed an informed consent form prior to participation, and the research was done in accordance with the Declaration of Helsinki (2008).

#### Apparatus and Stimuli

Participants were individually seated in dimly-lit sound-attenuated testing cabins, approximately 60 cm from a 22’ CRT screen (Iiyama MA203DT). The resolution was set to 1280 by 1024 pixels, at 16-bit color depth and 100 Hz refresh rate. Open Sesame 3.2 (Mathôt et al., 2012) with the Expyriment back-end (Krause & Lindemann, 2013) was used for trial preparation and data collection, running under the Microsoft Windows 7 operating system. Responses were collected with a Logitech Attack 3 joystick.

The background color was set to grey (RGB 128, 128, 128). Target stimuli were orientation gratings with a spatial frequency of 1.8 cycles/degree, and a standard deviation of .27°, presented within a circle spanning 2.1° of visual angle (Figure 1b, bottom left of the dashed line; colored targets shown to the bottom right of the dashed line were used in Experiment 2). The orientation of the lines was chosen randomly from any angle between 1 to 180°. Furthermore, the orientation difference between the first and second target was always at least 20 degrees in order to limit the similarity between targets (Figure 1c). The distractors/maskss (Figure 1b, top left of the dashed line; colored distractors shown to the top right of the dashedd line were used in Experiment 2) were scrambled, mosaic-patched orientation gratings. These were built as follows: First, 150 orientation gratings, which were identical to target stimuli in terms of low-level stimuli features, were generated. Using Adobe Photoshop, each of thee orientation gratings was cut into squares of 5 by 5 pixels, which were subsequently randomly displaced. Each of the 5 by 5 pixel-squared areas was also randomly rotated. Finally, a mosaic patch was added to the image. As a result, the low-level features of targets and distractors matched, but any kind of orientation information was minimized.

#### Procedure

Each of the first two blocks consisted of 16 practice trials. Each trial started with a fixation dot, lasting from 600 to 1000 ms (Figure 1e). After the fixation dot, T1 appeared at either 1.5° visual angle above or below of the fixation dot, and remained on screen for 90 ms. At the same time, a distractor appeared on the other side of the fixation dot. After 10 ms of blank inter-stimulus-interval (ISI), a second array consisting of two distractors was shown at the same location as the first array, for 90 ms. After a brief (60 ms) or long (610 ms) ISI, the second target array appeared, such that the lag or SOA was either 250 or 800 ms. T2 appeared at 1.5° from its edge to either the left or right side of the fixation dot, again for 90 ms. As in the first target array, a distractor was shown on the other side of the fixation dot. The second target array was also followed by a distractor array, which consisted of two distractors appearing at the left and right side locations. Target location pairs were randomized but evenly distributed within and across both target arrays. Thus, both target location and temporal onset were unpredictable, providing no information to direct attention to either dimension voluntarily (cf. Denison, Heeger et al., 2017). 500 ms after the offset of the last array, participants were prompted to indicate the orientation of target items in the correct temporal order with a joystick. In the practice trials only, feedback was provided. If the target report was within 20 degrees of the actual target orientation, a happy smiley was shown. If the response was more than 20 degrees away from the target orientation, an unhappy smiley was shown. There was no trial-based feedback in experimental trials. However, after each block, feedback about their average error on each of the two targets was provided to participants. There were 10 experimental blocks in total and there were 64 trials within each block. Participants were able to have a self-timed break between blocks. The experiment took approximately 75 minutes per participant.

#### Design and Analysis

As in virtually all AB designs, SOA between T1 and T2 was manipulated. There were two conditions in the experiment; a short SOA of 250 ms, and a long SOA of 800 ms, constituting a 2-factorial within-subjects experimental design. Initial analyses were conducted on reproduction error reflecting the absolute angular difference of target and response value. Error was assessed for both targets. As the AB is a consequence of the processing of the T1 on T2, performance on T2 was conditionalized on T1 (T2|T1), as is common in AB studies. Specifically, only trials in which T1 error was smaller than 22.5° were included. Using JASP version 0.9 (JASP Team, 2018), T1 and T2|T1 error across SOA were compared by means of Bayesian paired-sample T-tests. The resultant Bayes factors (BF) were interpreted according to Wetzels et al. (2011). BF_10_ values between 1 to 3 were considered as anecdotal evidence, 3 to 10 as substantial evidence, 10 to 30 as strong evidence, 30 to 100 as very strong evidence, and BF_10_ values above 100 as decisive evidence in favor of the alternative hypothesis in a given test. BF_10_ values between 0.33 to 1 were considered as anecdotal evidence, 0.1 to 0.33 as substantial evidence, 0.03 to 0.33 as strong evidence, 0.01 to 0.03 as very strong evidence, and BF_10_ values below 0.01 as decisive evidence in favor of the null hypothesis in a given test.

Apart from computing overall response error, the data from both targets were modeled by applying a standard mixture model in the CatContModel package (Version 0.8.0; Hardman, 2016) in R 3.6.3 (R Core Team, 2020), which computes estimates of guess rate and precision (Zhang and Luck, 2008; Bays et al., 2009). Guess rate (Pg) reflects the probability of guessing a target’s identity without evidence for any underlying target information, as is reflected in the height of the uniform distribution (Figure 1a). Precision (the inverse of σ) reflects the degree to which some target information was retained above and beyond guessing, as evidenced by the standard deviation of responses systematically approaching zero with increased precision. The guess rate and precision parameters allow for an assessment of the degree to which target perception occurs in an all-or-none, or in a gradual fashion, respectively. Thus, by applying the mixture model, underlying sources of error that contribute to overall task accuracy can be determined and analyzed.

#### Model-based Comparisons

In order to statistically infer the precense or absence of guess rate and precision effects, we used a model comparison approach, which consisted of two steps: (I) Model comparison, to identify and select the optimal model, with the least number of parameters, and (II) hypothesis testing to perform Bayesian hypothesis testing on guess rate and precision estimated by the best fitting model. In more detail, following Ricker and Hardman (2017) and Hardman, Vergauwe, and Ricker (2017), to evaluate how model parameters (Pg and σ) change across SOA conditions (step I), we applied hierarchical Bayesian estimation (as implemented by the CatContModel package), taking into account each participant as a sample of the population. First of all, a full model in which both Pg and σ vary between SOA conditions was calculated. Next, we generated reduced models by keeping either one or both of the parameters of the full model constant across conditions. We then estimated model parameters with full and reduced models, fitted the models to the data, and calculated the Watanabe-Akaike Information Criterion (WAIC) for each model. WAIC penalizes models on the number of free parameters in a modest manner and lower WAIC scores indicate better fit (Gelman et al., 2014). Parameter estimates and fit statistics were also calculated with the CatContModel package. In order to estimate the parameters, we ran 11000 iterations with the Bayesian Markov Chain Monte Carlo (MCMC) sampling technique. The first 1000 iterations were discarded as burn-in, and Metropolis-Hastings Tuning was applied. For each model, a total of 10000 MCMC iterations were thus used to estimate model parameters and fits. After having selected the best model, the remaining model parameters were compared between SOA conditions (step II), with Bayesian tests that are conceptually equivalent to ANOVAs (see Ricker & Hardman, 2017). For example, if keeping Pg constant across SOA conditions was the best model, then Bayesian comparisons were done only on the σ parameter across SOA conditions. The outcomes of both analysis steps were visualized with the “ggplot2” package (Wickham, 2016).

Finally, to confirm that true underlying models and parameters could be recovered successfully with our methods, we performed model recovery analyses (see Supplementary Material). Data were generated from the standard mixture model, with parameters and sample sizes matching the experiments of this study. In general, true models consistently had the lowest WAIC, validating our approach. WAIC scores were biased towards selecting more complex models, possibly due to common variance explained by both guess rate and standard deviation when the standard deviation is high. Nevertheless, guess rate and precision effects were not confused: Fits were biased to models where both parameters varied, not to models where the other parameter varied. For instance, when the true model only contained precision effects, the full model containing both effects fitted relatively well, but the model with only guess rate effects fitted relatively poorly. These results suggest that our model-based approach can succesfully separate effects of discrete and gradual loss of awareness in our experimental paradigm.

#### Data availability

The study was not preregistered, but in order to increase replicability, and to provide scientific transparency, the raw data, model outcomes, the analysis scripts, and an example of the experimental tasks were uploaded in a publication package, with the unique identifier x5dru, to the Open Science Framework (https://osf.io/x5dru).

### Results

#### Error

On T1 error, Bayesian T-tests showed decisive evidence in favor of the alternative hypothesis, indicating an effect of SOA (BF_10_ = 579.7). T1 error was 26.1° at short SOA and decreased to 23.5° at long SOA (Table 1; see Supplementary Material for T1-related figures for this and all other experiments). On T2|T1 error (Figure 2a, b), there was also decisive evidence in favor of the alternative hypothesis for the existence of an effect of SOA (BF_10_ > 1000). T2|T1 error averaged 32.9° at short SOA, compared to 26.5° at long SOA.

**Figure 2.**
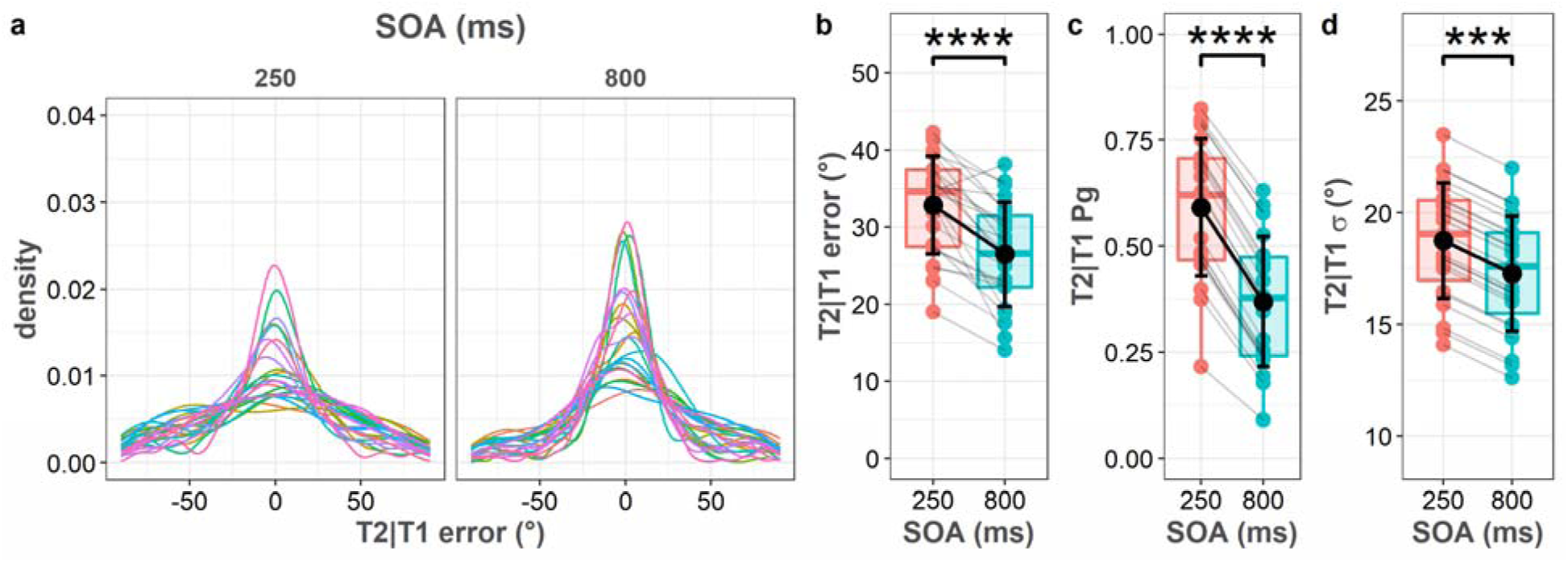
T2|T1 performance in Experiment 1A. **a** Probability density plot of the T2|T1 error distribution of each subject for each SOA condition. Each color indicates a subject. **b** T2|T1 error for short and long SOA conditions (ms). Colored dots represent individual data, black dots reflect averages, and error bars represent 95% confidence intervals. Grey lines connect each subject in short and long SOA conditions. Whisker plots show median and quartile values. The orange color shows the short SOA condition and the blue color shows the long SOA condition. * indicates anecdotal evidence against the null hypothesis, ** indicates substantial evidence against the null hypothesis, *** indicates strong evidence, and **** means decisive evidence. **c** T2|T1 Pg based on the best model predictions. **d** T2|T1 σ based on the best model predictions.

**Table 1.** T1 and T2|T2 error, Pg, and σ estimates from the best fitting model across conditions

|  |  | T1 |  |  |  |  |  | T2\|T1 |  |  |  |  |  |
| --- | --- | --- | --- | --- | --- | --- | --- | --- | --- | --- | --- | --- | --- |
| | | error (°) | | $P_g$ | | $\sigma$ (°) | | error (°) | | $P_g$ | | $\sigma$ (°) | |
| | | $\bar{x}$ | SEM | $\bar{x}$ | SEM | $\bar{x}$ | SEM | $\bar{x}$ | SEM | $\bar{x}$ | SEM | $\bar{x}$ | SEM |
| Exp 1a | Short SOA | 26.12 | 1.95 | 0.42 | 0.05 | 13.96 | 0.5 | 32.85 | 1.33 | 0.59 | 0.03 | 18.75 | 0.54 |
|  | Long SOA | 23.45 | 1.85 | 0.35 | 0.05 | 13.96 | 0.5 | 26.46 | 1.42 | 0.37 | 0.03 | 17.28 | 0.54 |
| Exp 1b | Short SOA | 23.64 | 1.88 | 0.35 | 0.05 | 13.7 | 0.35 | 33.12 | 1.28 | 0.62 | 0.04 | 16.47 | 0.35 |
|  | Long SOA | 21.5 | 1.75 | 0.29 | 0.05 | 13.7 | 0.35 | 23.83 | 1.39 | 0.34 | 0.04 | 14.6 | 0.35 |
| Exp 2a | Short SOA | 17.85 | 2.14 | 0.08 | 0.03 | 16.36 | 0.53 | 20.27 | 1.58 | 0.08 | 0.02 | 18.54 | 0.59 |
|  | Long SOA | 16.69 | 2.1 | 0.08 | 0.03 | 15.57 | 0.53 | 18.59 | 1.45 | 0.08 | 0.02 | 17.13 | 0.59 |
| Exp 2b | Short SOA | 20.74 | 0.88 | 0.07 | 0.01 | 19.63 | 0.47 | 26.53 | 1.72 | 0.14 | 0.02 | 21.61 | 0.41 |
|  | Long SOA | 18.99 | 0.75 | 0.05 | 0.01 | 19.22 | 0.47 | 20.7 | 0.94 | 0.07 | 0.01 | 20.03 | 0.41 |
| Exp 3 | Short SOA | 19.79 | 1.66 | 0.23 | 0.04 | 14.76 | 0.68 | 32.77 | 1.84 | 0.6 | 0.05 | 17.05 | 0.81 |
|  | Long SOA | 17.7 | 1.61 | 0.18 | 0.03 | 14.06 | 0.68 | 20.29 | 1.39 | 0.16 | 0.03 | 17.05 | 0.81 |
| Exp 4 | Short-Same | 22.02 | 1.27 | 0.3 | 0.04 | 14.05 | 0.31 | 28.09 | 1.29 | 0.46 | 0.03 | 15.87 | 0.32 |
|  | Short-Diff | 19.99 | 1.06 | 0.24 | 0.03 | 14.05 | 0.31 | 28.48 | 1.28 | 0.46 | 0.03 | 15.87 | 0.32 |
|  | Long-Same | 20.53 | 1.12 | 0.25 | 0.03 | 14.05 | 0.31 | 22.2 | 1.08 | 0.3 | 0.03 | 14.93 | 0.32 |
|  | Long-Diff | 19.57 | 1.12 | 0.23 | 0.03 | 14.05 | 0.31 | 22.61 | 1.25 | 0.3 | 0.03 | 14.93 | 0.32 |
| Exp 5 | Short SOA | 21.18 | 1.44 | 0.23 | 0.04 | 15.67 | 0.57 | 29.8 | 1.92 | 0.51 | 0.05 | 16.02 | 0.38 |
|  | Long SOA | 20.58 | 1.57 | 0.23 | 0.04 | 15.11 | 0.57 | 25.35 | 1.57 | 0.37 | 0.05 | 16.02 | 0.38 |

Exp = experiment, $P_g$ = probability of guesses, $\sigma$ = memory (im-)precision, $\bar{x}$ = mean, SEM = standard error of the mean, Short-Same = Short SOA and same location condition, Short-Diff = Short SOA and different location condition, Long-Same = Long SOA and same location condition, Long-Diff = Long SOA and different location condition

#### Model Comparisons

For T1, the best model was the constant σ across SOA model (Table 2). For T2|T1, the best model was the full model, indicating that both Pg and σ differed between SOA conditions (Figure 2c, d). The inclusion of σ in the T2|T1 model is the first evidence of gradual perceptual awareness during the AB.

**Table 2.** WAIC for All Tested Models in Experiment 1.

| Model | WAIC | T1 | WAIC | T2\|T1 |
| --- | --- | --- | --- | --- |
|  |  | Difference From the Best Model |  | Difference From the Best Model |
| Full Model | 158374.80 | 1.08 | **102280.65** | **0.00** |
| Constant $\sigma$ across SOA | **158373.70** | **0.00** | 102283.19 | 2.54 |
| Constant $P_g$ across SOA | 158414.40 | 40.67 | 102341.78 | 61.13 |
| Constant $\sigma$ and $P_g$ across SOA | 158422.70 | 48.96 | 102517.09 | 236.44 |

Parameters were estimated freely in the full model. In the constant $g$ model, $P_g$ was invariant across SOA conditions, and $\sigma$ was freely estimated for both short and long SOA conditions. In the constant $\sigma$ model, $\sigma$ was invariant, and $P_g$ was estimated for short and long SOA conditions. In the constant $\sigma$ and $P_g$ model, a single $P_g$ and $\sigma$ were estimated across SOA conditions.

#### Hypothesis Testing

Constant σ across SOA was the best model for T1 error. Therefore, we tested the main effect of SOA on T1 Pg. Bayesian tests showed decisive evidence in favor of the main effect of SOA on T1 Pg (BF_10_ > 1000). T1 Pg was 0.42 at short SOA and decreased to 0.35 at long SOA. Given that the full model was the best model explaining T2|T1 error, we tested the main effect of SOA on both T2|T1 Pg and σ. Test results revealed decisive evidence in favor of the effect of SOA on Pg and substantial evidence on σ (BF_10_ > 1000 and BF_10_ = 8.72, respectively). T2|T1 Pg was 0.59 at short SOA and decreased to 0.37 at long SOA (Figure 2c). T2|T1 σ averaged 18.8° at short SOA compared to 17.3° at long SOA (Figure 2d).

### Discussion

It seems that in the DT paradigm, perceptual awareness during the AB is not only all-or-none, as expressed by a guess rate effect, but is also of a gradual nature, producing a precision effect as well. In other words, the representation of T2 suffers from T1 processing, but the representation is not always completely lost, with some evidence that T1 was affected similarly (cf. Hommel & Akyürek, 2005). This finding stands in contrast to previous findings by Asplund et al. (2014) and Tang et al. (2020), who obtained only all-or-none patterns in their continuous report RSVP tasks. The current outcome thus suggests that there might be something fundamentally different in the nature of perceptual awareness during the AB as observed in DT and RSVP tasks. The most obvious factor might be that of space; target locations vary in the DT paradigm, whereas they typically do not in RSVP. As a first step towards assessing the possible role of space in gradual awareness during the AB, a new DT task was implemented in Experiment 1B, in which target locations were made (even) more variable.

## Experiment 1B

The experiment was identical to Experiment 1A with only one difference: T1 was shown on either the horizontal or vertical pair of locations, instead of on the vertical pair only, while T2 was always on the other pair. If gradual awareness is a function of the number of possible target locations, the effect of SOA on T2 report precision should be increased in the current task.

### Method

### Participants

Twenty-six new students participated in the study in exchange for course credits or monetary reward. One participant did not complete the experiment, and was therefore excluded from the analysis. Four further participants were excluded because their error rate was at chance level in the long SOA condition, so that 21 participants remained (13 females, 8 males; mean age = 19.8, range = 18-24).

#### Procedure and design

Experiment 1B was identical to Experiment 1A, but with two exceptions. First, the duration of the fixation dot was random and varied from 300 ms to 500 ms in order to decrease the duration of the experiment, and second, T1 could appear on either the vertical or horizontal pair of locations. If T1 appeared on the vertical pair (as in Experiment 1A), T2 appeared on the horizontal pair, and vice versa.

### Results

#### Error

Bayesian paired-sample T-tests revealed strong evidence for the effect of SOA on T1 error, and decisive evidence on T2|T1 error (BF_10_ = 14.8; BF_10_ > 1000, respectively). T1 error averaged 23.6° in the short SOA condition and 21.5° in the long SOA condition (Table 1). Moreover, T2|T1 error (Figure 3a, b) averaged 33.1° at short SOA and decreased to 23.8° at long SOA.

**Figure 3.**
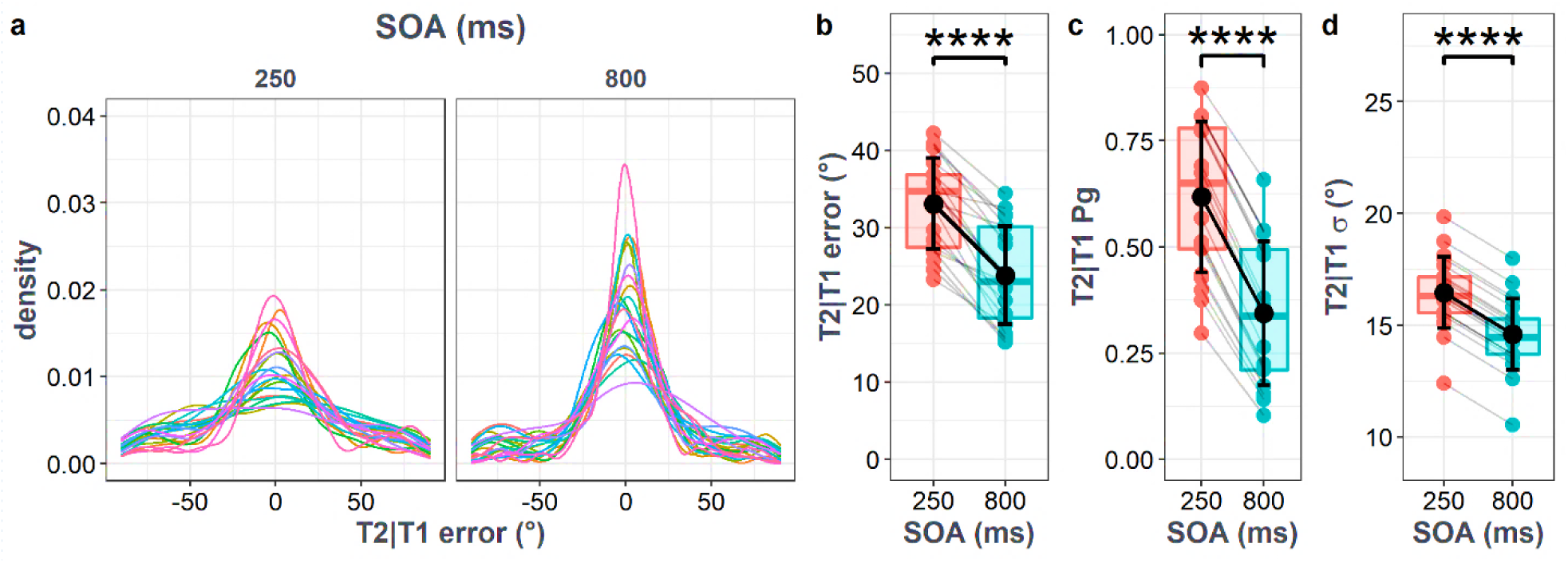
T2|T1 performance in Experiment 1B. **a** Probability density plot of the T2|T1 error distribution of each subject for each SOA condition. **b** T2|T1 error, **c** T2|T1 Pg, **d** T2|T1 σ for short and long SOA conditions. Figure conventions follow Figure 2.

#### Model Comparisons

On T1, the constant σ across SOA model was the best, and on T2|T1 it was the full model, in which both P_g_ and σ vary, depending on the SOA (Table 3).

**Table 3.** WAIC for All Tested Models in Experiment 1B

| Model | WAIC | T1 | WAIC | T2\|T1 |
| --- | --- | --- | --- | --- |
|  |  | Difference From the Best Model |  | Difference From the Best Model |
| Full Model | 143968.60 | 2.06 | **100068.30** | **0.00** |
| Constant $\sigma$ across SOA | **143966.60** | **0.00** | 100077.72 | 9.42 |
| Constant $Pg$ across SOA | 143994.10 | 27.57 | 100196.33 | 128.03 |
| Constant $\sigma$ and $Pg$ across SOA | 144000.90 | 34.32 | 100489.50 | 421.20 |

#### Hypothesis Testing

There was decisive evidence supporting the effect of SOA on T1 Pg (BF_10_ >> 1000). T1 Pg averaged 0.35 at short SOA and decreased to 0.29 at long SOA. Furthermore, decisive evidence was observed in favor of the main effect of SOA on T2|T1 Pg and T2|T1 σσ (BF_10_ > 1000; BF_10_ = 153.9, respectively; Figure 3c, d). T2|T1 Pg averaged 0.62 at short SOA andd 0.34 at long SOA. Lastly, T2|T1 σ averaged 16.5° at short SOA and decreased to 14.6° at longg SOA.

### Discussion

The results of the current DT task, in which target locations were varied across the horizontal and vertical stimulus location pairs, were very similar to those of Experiment 1A. The principal finding of an effect of SOA on target report precision was replicated. One might even argue that Experiment 1B provided stronger evidence than Experiment 1A, since the WAIC values for the model with a variable guess rate only were higher in the former. Overall, the increased variability of target locations did not seem to result in a very different outcome qualitatively, suggesting that distributing attention across more than a single location is already sufficient to induce gradual awareness during the AB. The role of space in gradual awareness will be further explored in subsequent experiments. First, however, it is necessary to assess whether our choice of stimuli (orientation gratings) might have confounded the results.

## Experiment 2A

Previous studies that reported discrete, rather than gradual target awareness during the AB have used color stimuli (e.g., Asplund et al., 2014). To rule out the possibility that gradual awareness during the AB might be specific to orientation stimuli, Experiment 2A was set up with color instead of orientation stimuli, to replicate and generalize the results of Experiment 1A.

### Method

#### Participants

26 new students (22 females, 4 males) participated in the study in exchange for course credits (mean age = 19.9, range = 18-28).

#### Apparatus, Stimuli, and Procedure

Experiment 2A was identical to Experiment 1 with the following changes (see Figure 1d, f). Target stimuli were circles filled with colors from the HSL color spectrum. Saturation was set to 100% and luminance was set to 50%. Hue was chosen randomly from the HSL color wheel. The task was to reproduce the exact color of the targets from a color wheel and responses were collected with a standard mouse. In view of the robust effects, and in order to counteract possible fatigue, there were now 8 blocks with 64 trials each, which meant that the experiment lasted approximately one hour.

### Results

#### Error

Very strong evidence was observed for an influence of SOA on T1 and T2|T1 error (BF_10_ = 64.3 and BF _10_ = 61.5, respectively). The effect size appeared to be modest: T1 error was 17.9° at short SOA and decreased to 16.7° at long SOA (Table 1). T2|T1 error averaged 20.3° at short SOA and decreased to 18.6° at long SOA (Figure 4a, b).

**Figure 4.**
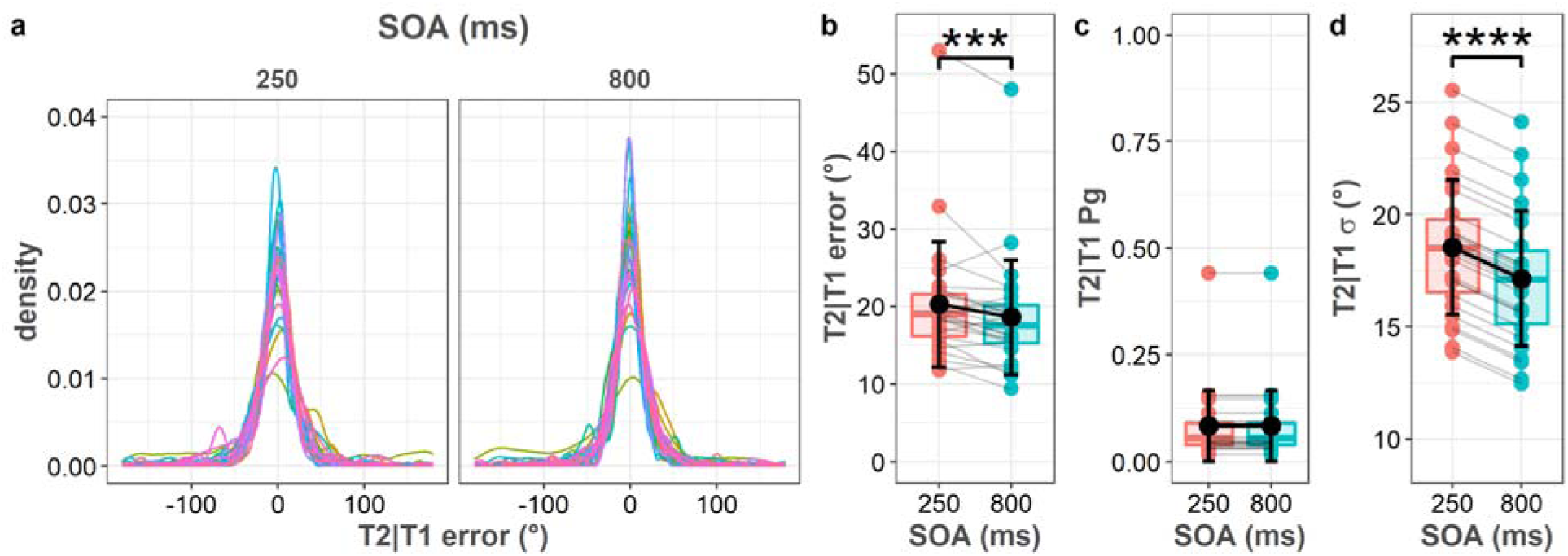
T2|T1 performance in Experiment 2A. **a** Probability density plot of the T2|T1 error distribution of each subject for each SOA condition. **b** T2|T1 error, **c** T2|T1 Pg, **d** T2|T1 σ for short and long SOA conditions. Figure conventions follow Figure 2.

#### Model Comparisons

Model comparisons revealed that for T1 the best model was the constant P_g_model (Table 4). For T2|T1, the best model was the constant P_g_ model, indicating that only precision differed depending on the SOA condition (Figure 4c, d). Thus, as in Experiment 1, the model comparisons provided evidence for gradual perceptual awareness of blinked targets.

**Table 4.**
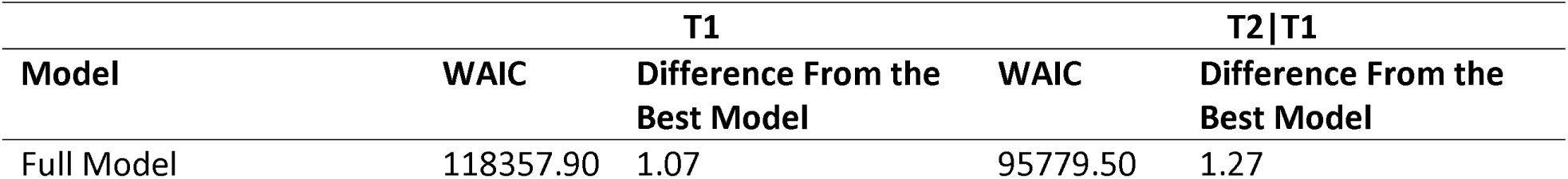
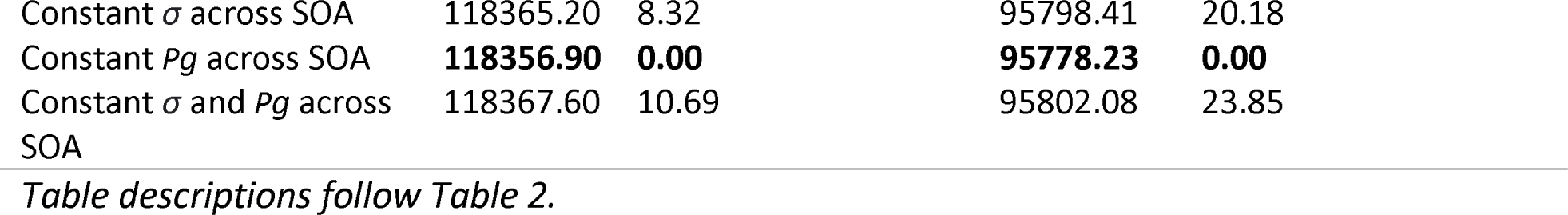
WAIC for All Tested Models in Experiment 2A.

| Model | WAIC | T1 | WAIC | T2\|T1 |
| --- | --- | --- | --- | --- |
|  |  | Difference From the Best Model |  | Difference From the Best Model |
| Full Model | 118357.90 | 1.07 | 95779.50 | 1.27 |
| Constant $\sigma$ across SOA | 118365.20 | 8.32 | 95798.41 | 20.18 |
| Constant $Pg$ across SOA | **118356.90** | **0.00** | **95778.23** | **0.00** |
| Constant $\sigma$ and $Pg$ across SOA | 118367.60 | 10.69 | 95802.08 | 23.85 |

#### Hypothesis Testing

There was very strong evidence supporting the effect of SOA on T1 σ (BF_10_ = 56.6). T1 σ averaged 16.4° at short SOA and 15.6° at long SOA (Table 1). Additionally, decisive evidence in favor of the effect of SOA on T2|T1 σ existed (BF_10_ > 1000), again replicating the findings of Experiment 1. T2|T1 σ averaged 17.1° at long SOA compared to 18.5° at short SOA (Figure 4d).

### Discussion

The results showed a compelling effect of SOA on T2 report precision, coupled with an absence of any effect of guess rate. Again, there was evidence against a guess-only account also, since the WAIC value for this model (constant σ) was high. It thus seems that the orientation stimuli used in Experiment 1 were not a causal factor in obtaining a precision effect. If anything, the same task with color stimuli showed even stronger support for gradual awareness during the AB.

## Experiment 2B

Experiment 2A replicated the results of Experiment 1. However, the color stimuli used in the former experiment did result in high task performance and small blink magnitude. To ensure that this high level of task performance did not confound the findings, task difficulty was increased in Experiment 2B by reducing the salience of the targets, as follows: First, we used colored distractors instead of black and white distractors. Even though it has been shown that the AB occurs even with minimal 4-dot masking (Dell’Acqua, Pascali, Jolicœur, & Sessa, 2003), and even without any T1 mask (Visser, 2007), a stronger mask is still likely to increase blink magnitude. Second, we used CIELAB colors as described in Zhang and Luck (2008), which are more perceptually uniform than HSL colors.

### Method

#### Participants

27 new students participated in the study in exchange for course credits (17 females, 10 males; mean age = 19.33, range = 18-22).

#### Stimuli, Apparatus, and Procedure

Experiment 2B was identical to 2A with the following changes: As in Experiment 1, there were ten blocks of experimental trials and each block consisted of 64 trials. Both targets and distractors were now rendered in CIELAB color space (see Figure 1b). The procedure to generate the distractors was similar to that in Experiment 2A, but now used six random colors from the CIELAB color wheel. First, a circle with the same size as the targets was drawn, sliced into six equal pieces. Each piece was filled with one of the six random colors. The background was filled with gray. Afterwards, each colored distractor was cut into squares of 5 by 5 pixels, randomly rotated and displaced, followed by a mosaic patch. There were 150 different colored distractors in total. Results Error: Decisive evidence was observed of the effect of SOA on T1 error (B_10_ = 391.5). T1 error was 20.7° at short SOA and decreased to 19.0° at long SOA (Table 1). Decisive evidence was also obtained for an SOA effect on T2|T1 error (BF_10_ > 1000). T2|T1 error was 26.5° at short SOA, and 20.7° at long SOA (Figure 5a, b), representing a substantial effect size.

**Figure 5.**
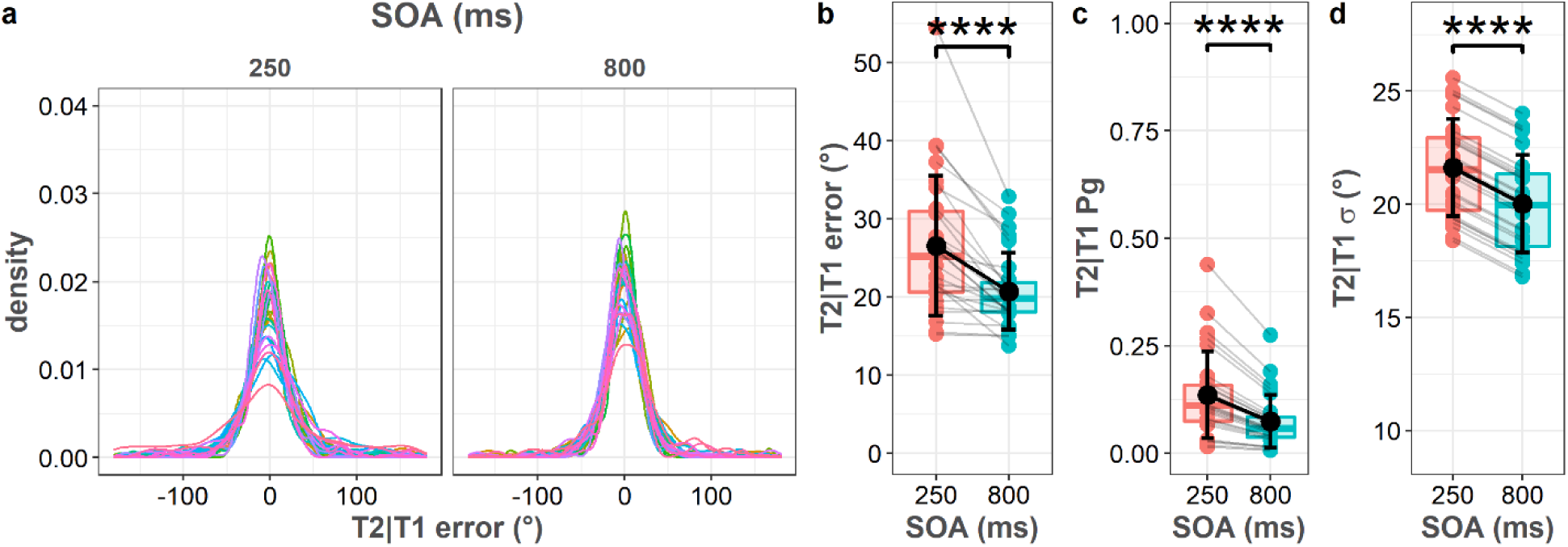
T2|T1 performance in Experiment 2B. **a** Probability density plot of the T2|T1 error distribution of each subject for each SOA condition. **b** T2|T1 error, **c** T2|T1 Pg, **d** T2|T1 σ for short and long SOA conditions. Figure conventions follow Figure 2.

#### Model Comparisons

For both T1 and T2|T1, the best model was the full model (Table 5). These model comparisons indicate that both T2|T1 Pg and σ differed between SOA conditions, again suggesting there was gradual awareness of blinked targets (Figure 5c, d).

**Table 5.**
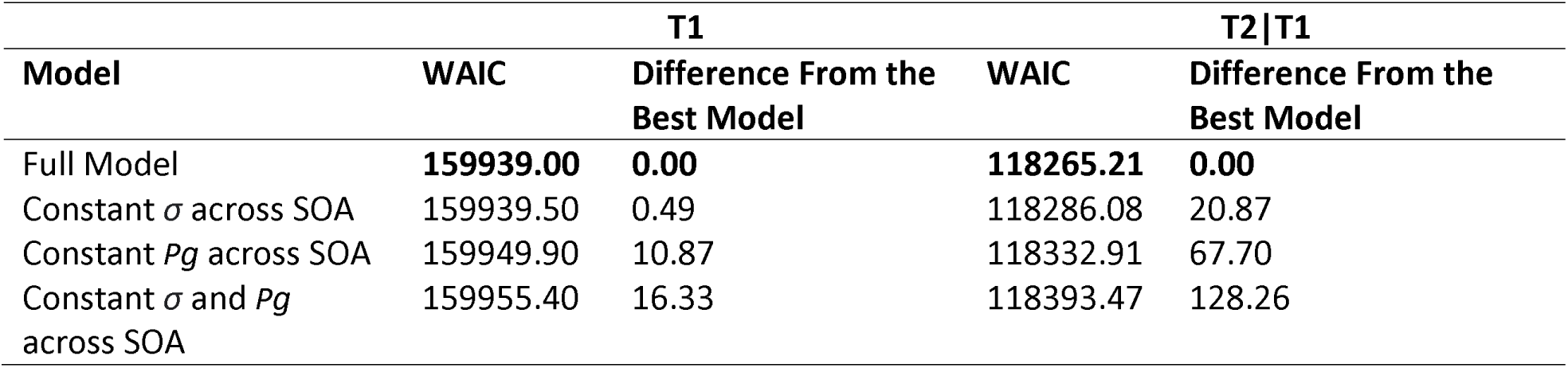
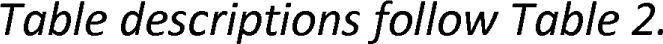
WAIC for All Tested Models in Experiment 2B.

#### Hypothesis Testing

Further to the model comparisons, Bayesian test results revealed very strong evidence in favor of the alternative hypothesis that SOA influenced T1 Pg, (BF_10_ = 72.4). T1 Pg averaged 0.07 in the short SOA condition and decreased to 0.05 in the long SOA condition. Also, there was anecdotal evidence in favor of the effect of SOA on T1 σ (BF_10_ = 1.1). T1 σ was 19.6° at short SOA and 19.2° at long SOA. There was decisive evidence in favor of the alternative hypothesis that SOA influenced T2|T1 Pg as well as T2|T1 σ (both BF_10_ > 1000). T2|T1 Pg (Figure 5c) was 0.14 at short SOA, and 0.07 at long SOA. T2|T1 σ (Figure 5d) was 21.6° at short SOA, and 20.0° at long SOA.

### Discussion

The results of Experiment 2B largely replicated those of Experiments 1A, 1B, and 2A, providing further evidence that at short SOA, the AB resulted in reduced T2|T1 precision, giving rise to gradually impaired target awareness in the DT paradigm. In all three experiments, we observed consistent precision effects, in tow with guess rate effects in Experiment 1 and 2B. Altogether, the evidence confirms that the nature of the AB in the DT paradigm is to a considerable extent gradual, and not only discrete; the AB caused a variable loss of quality in the representation of both orientation and color targets.

## Experiment 3

Experiment 1 and 2 showed a clear dissociation with the existing literature, in which perceptual awareness during the AB has been characterized exclusively as an all-or-none phenomenon (Asplund et al., 2014; Sergent & Dehaene, 2004; Sergent et al., 2005; Tang et al., 2020). Given that this discrepancy does not appear to be due to the nature of the stimuli (i.e., orientation or color), the results suggest that there might be a fundamental, underlying difference in the performance deficits observed in DT and RSVP tasks, even though both paradigms elicit the typical AB. The gradual nature of the AB as presently observed could be a result of task differences between DT and RSVP paradigms that change the way that temporal attention operates. One of the most conspicuous differences between the DT and RSVP tasks is the inherently spatial nature of the former. Before investigating this issue, it is essential to verify first that attentional allocation is indeed different in the DT and RSVP tasks. Experiment 3 was therefore aimed at replicating previous reports of pure guess rate effects in RSVP.

### Method

#### Participants

26 new students participated in the study for course credit. Two of them were excluded from the analysis, following the same criteria as in Experiment 1, leaving 15 females and 9 males (mean age = 19.6, range = 18-23).

#### Stimuli, Apparatus, and Procedure

Experiment 3 was identical to Experiment 1 with the following changes. A classical RSVP design was applied (Figure 6a). Each trial started with a fixation dot ranging from 300 to 500 ms. After the fixation dot, an RSVP consisting of 18 items started (2 targets and 16 distractors). Targets and distractors were identical to Experiment 1. Each item in the stream was shown in the center of the screen for 90 ms and succeeding items in the RSVP were separated with 10 ms ISI. T1 was the 5^th^ to 8^th^ item in the stream, which was randomized but evenly distributed. The number of distractors between targets differed depending on the experimental manipulation. There were two distractors (SOA = 300 ms) between T1 and T2 in the short SOA condition and 7 distractors (SOA = 800 ms) in the long SOA condition. After the stream ended there was a 500 ms interval before the response prompts appeared. As before, the task was to reproduce the orientations of the targets in the correct temporal order. There were 8 experimental blocks, and each block consisted of 60 trials.

**Figure 6.**
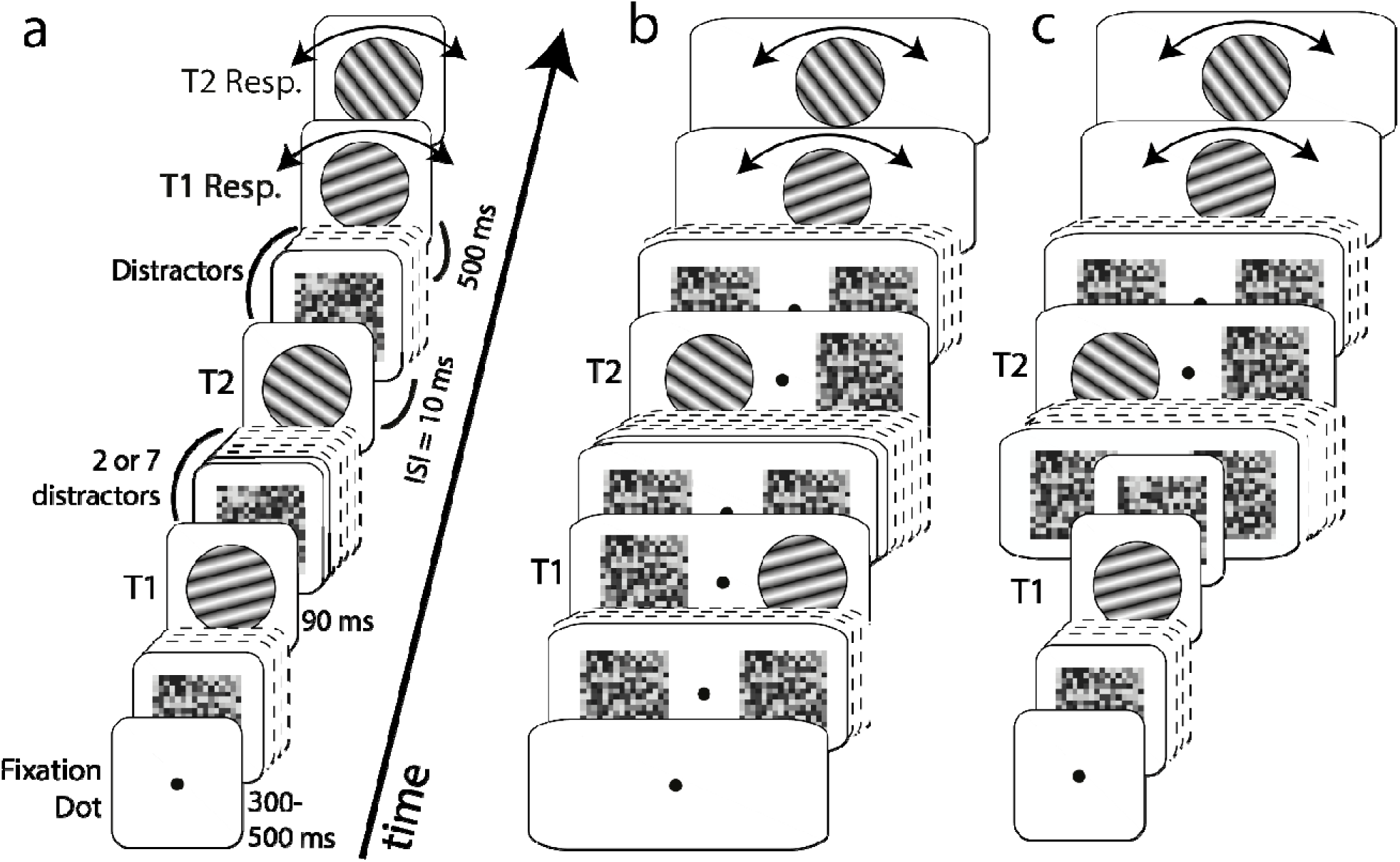
The experimental tasks used in Experiment 3, 4, and 5. The actual size of the stimulli was identical to Experiment 1 and 2, as was the distance between the center and target/distractor locations. T1 Resp means T1 response prompt and T2 Resp means T2 response prompt. **a** Trial schematic of Experiment 3. Dashed lines indicate a variable number of distractors. **b** Trial schematic of Experiment 4. A different target location trial is shown. Stimulus timing was identical to Experiment 3. **c** Trial schematic of Experiment 5. T1 is presented within aa single central stream. After the T1+1 distractor, the central stream disappeared, and two new lateral streams started. T2 then appeared either in the right or in the left side stream.

### Results

#### Error

Decisive evidence was observed for the effect of SOA on T1 error (BF_10_> 1000). T1 error (Table 1) was 19.8° at short SOA and 17.7° at long SOA. Decisive evidence was also obtained for an SOA effect on T2|T1 error (BF_10_ > 1000). T2|T1 error (Figure 7a, b) was 32.8° at short SOA, and 20.3° at long SOA. <u>Model Comparisons:</u> The full model was the best model for T1. Constant σ across SOA was the best model for the effect of SOA on T2|T1 (Table 6). This finding showed evidence in favor of discrete awareness (Figure 7c, d), which is in line with existing accounts of the AB (Asplund et al., 2014; Tang et al., 2020; Dehaene et al., 2003; Sergent et al., 2005).

**Figure 7.**
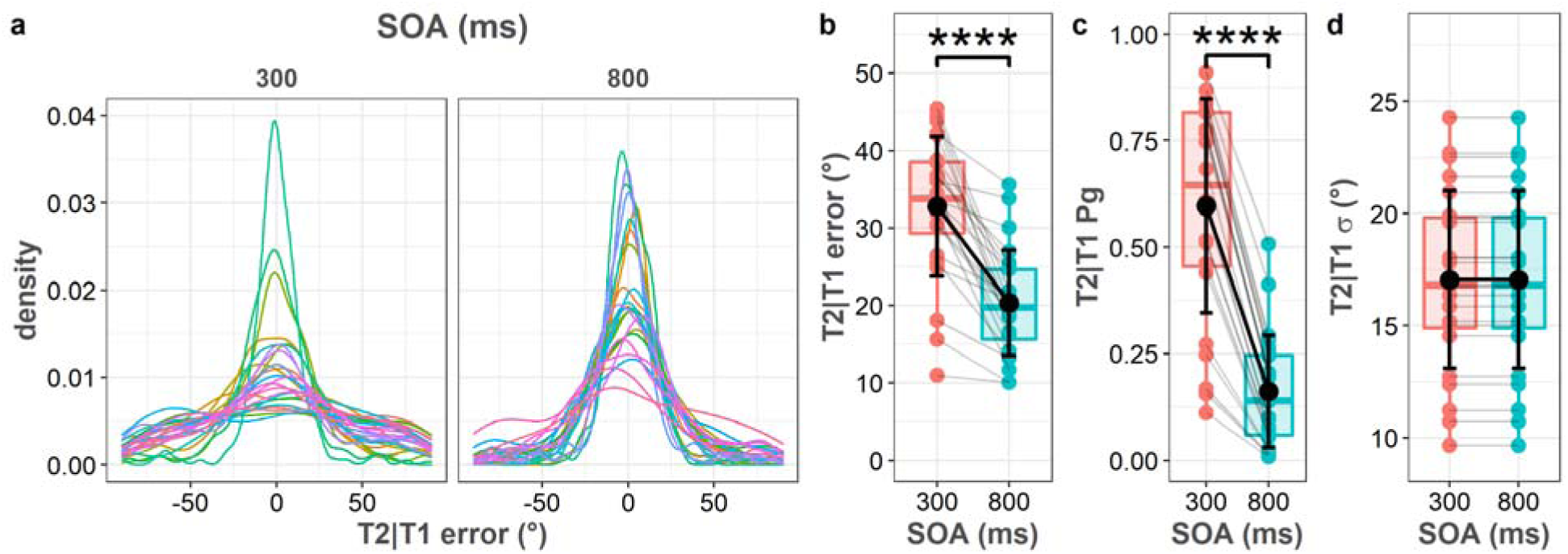
T2|T1 performance in Experiment 3. **a** Probability density plot of the T2|T1 error distribution of each subject for each SOA condition. **b** T2|T1 error, **c** T2|T1 Pg, **d** T2|T1 σ for short and long SOA conditions. Figure conventions follow Figure 2.

**Table 6.** WAIC for All Tested Models in Experiment 3.

| Model | WAIC | T1 | WAIC | T2\|T1 |
| --- | --- | --- | --- | --- |
|  |  | Difference From the Best Model |  | Difference From the Best Model |
| Full Model | **117418.50** | **0.00** | 90510.12 | 3.06 |
| Constant $\sigma$ across SOA | 117426.60 | 8.08 | **90507.06** | **0.00** |
| Constant $Pg$ across SOA | 117435.80 | 17.29 | 90947.20 | 440.14 |
| Constant $\sigma$ and $Pg$ across SOA | 117458.50 | 40.00 | 91278.28 | 771.22 |

#### Hypothesis Testing

Decisive evidence for an effect of SOA on T1 Pg was observed (BF_10_= 255.3). T1 Pg (Table 1) was 0.23 at short SOA, and decreased slightly to 0.18 at long SOA. There was very strong evidence supporting the effect of SOA on T1 σ (BF_10_= 54.4). T1 σ was 14.8° at short SOA and 14.1° at long SOA. There was also decisive evidence in favor of the effect of SOA on T2|T1 Pg (BF_10_> 1000). T2|T1 Pg (Figure 7c) was 0.60 at short SOA, compared to 0.16 at long SOA.

### Discussion

The analyses showed that perceptual awareness was discrete in the current RSVP task, and thereby replicated the findings of Asplund et al. (2014) and Tang et al. (2020) for line orientations. Thus, in contrast to the DT tasks in Experiment 1A, 1B, 2A and 2B, the results of the current RSVP task were in line with discrete models of conscious awareness, such as the Global Neuronal Workspace theory (Dehaene et al., 2003), as well as major models of the AB, such as the eSTST theory (Wyble et al., 2009), which holds that T2 cannot bind to a token if attention is sufficiently suppressed during the AB window, depending on the strength of its representation, and is consequently lost altogether.

## Experiment 4

The results of the DT tasks in Experiment 1 and 2 contrasted with those of the RSVP task in Experiment 3, showing two faces of awareness during the AB: Either partially gradual, or fully discrete. Why did the nature of awareness change between these tasks? Possibly the most conspicuous difference between the cognitive processing required in DT and RSVP paradigms is that locations change in the former paradigm, but not in the latter. Spatial locations are known to be special when it comes to attentional processing. For instance, a location-specific cue will modulate the early P1 and N1 components of the event-related potential (Hillyard & Münte, 1984; Luck et al., 1994), while a feature-based cue has a noticeably later locus (Anllo-Vento & Hillyard, 1996; Hillyard & Münte, 1984). In the context of temporal attention, it has long been known that the Lag 1 sparing phenomenon that often accompanies the AB is prevented whenever a spatial switch occurs (Visser et al., 1999a). It is thus conceivable that a spatial switch may similarly change the way in which awareness is gated during the AB. Thus, to investigate the nature of awareness during the AB when targets occur in more than a single spatial location, a second RSVP stream was added in Experiment 4. If having a single spatial location is the critical factor in producing a predominantly all-or-none AB effect, then this should no longer be the case in the present experiment, and a gradual (i.e., precision) effect should be observed.

### Method

#### Participants

Thirty-two new students participated in the experiment in exchange for course credits or monetary compensation of 12 Euros. Eight of them were excluded from the analysis because there were less than 50 trials in at least one of the experimental conditions, so that 24 remained in the final sample (8 females, 16 males; mean age = 19.9, range = 18-23).

#### Stimuli, Apparatus, and Procedure

Experiment 4 was identical to Experiment 3 with the following changes. The fixation dot remained at the center of the screen from the start of the trial until the end to help participants to keep their gaze at the center. After the first fixation period (300 to 500 ms), two streams on the right and left side of the fixation dot started, at a lateral distance of 1.5° degrees of visual angle. Participants were instructed to keep fixating the central dot. T1 appeared in either the left or right stream, and again as the 5^th^ to 8^th^ item. The spatial location of T1 was randomized but evenly distributed. Apart from the usual lag between T1 and T2 (300 and 800 ms), the spatial location of T2 was also manipulated. Target locations were identical in the same location condition (e.g., T1 on the left, and T2 also on the left), and they were different in the different location condition (e.g., T1 on the right and T2 on the left, as shown in Figure 6b).

#### Design and Analysis

A 2 (SOA; short or long) by 2 (Location; same or different) design was used to investigate how spatial target locations influence identification performance. A Bayesian Repeated Measures ANOVA was run to test the main effect of SOA, Location, and their interaction on T1 and T2|T1 error. An inclusion Bayes factor, based on matched models (BF_inc-10_) was used instead of Bayesian model comparisons for the sake of simplicity (see Mathôt, 2017). Corrections were done for further post-hoc tests, by fixing prior probability to .5 (Westfall, Johnson, & Utts, 1997). A classical Bayesian Repeated Measures model comparison with multiple independent variables is provided in the Supplementary Material.

### Results

#### Error

Bayesian repeated-measures ANOVA results showed anecdotal evidence in favor of the alternative hypothesis that SOA affects T1 error (BF _inc-10_ = 2.7), but there was no evidence that Location influenced T1 error (BF _inc-10_ = .5). Nevertheless, there was decisive evidence for the interaction of SOA and Location on T1 error (BF _inc-10_ = 370.2). T1 error (Table 1) averaged 21.0° at short SOA and decreased to 20.1° at long SOA (BF_10_ = 1.7). T1 error was relatively high in the short SOA and same location condition, averaging 22.0°, while T1 error averaged 20.0° in the short SOA and different location condition (BF_10_= 22.1), and 20.5° in the long SOA and same location condition (BF_10_= 9.6). There was anecdotal evidence that T1 error in the latter condition also differed from that in the long SOA and different location condition (19.6°, BF_10_- = 2.3). T1 error in the short SOA and different location condition did not differ from that in the long SOA and different location condition (BF_10_= .4). On T2|T1 error (Figure 8a, b), there was decisive evidence in favor of the main effect of SOA (BF _inc-10_ > 1000). T2|T1 error averaged 28.3° at short SOA and 22.4° at long SOA (BF_10_ > 1000). There was substantial evidence against the effect of Location and against the interaction of SOA and Location on T2|T1 error (BF _inc-10_ = .3; BF _inc-10_ = .2, respectively).

**Figure 8.**
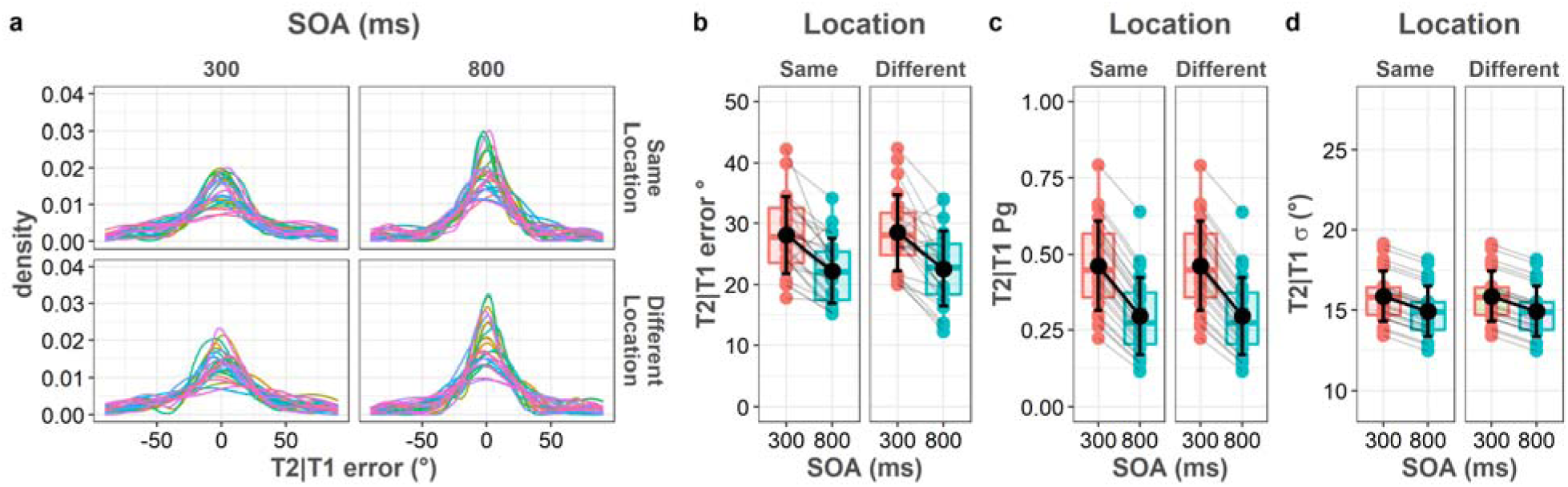
T2|T1 performance in Experiment 4. **a** Probability density plot of the T2|T1 error distribution of each subject for each SOA and Location condition. **b** T2|T1 error, **c** T2|T1 Pg, **d** T2|T1 σ for each SOA and Location condition. Figure conventions follow Figure 2.

#### Model Comparisons

We compared models across the main effects of SOA and Location as well as their interaction (Table 7). Similar to Experiment 1A and 1B, the best model for T1 had constant σ across SOA and Location. For T2|T1, the best model had constant σ and Pg across Location (Figure 8c, d).

**Table 7.**
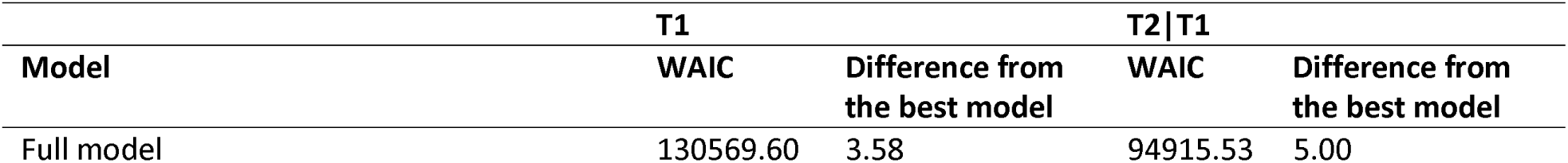
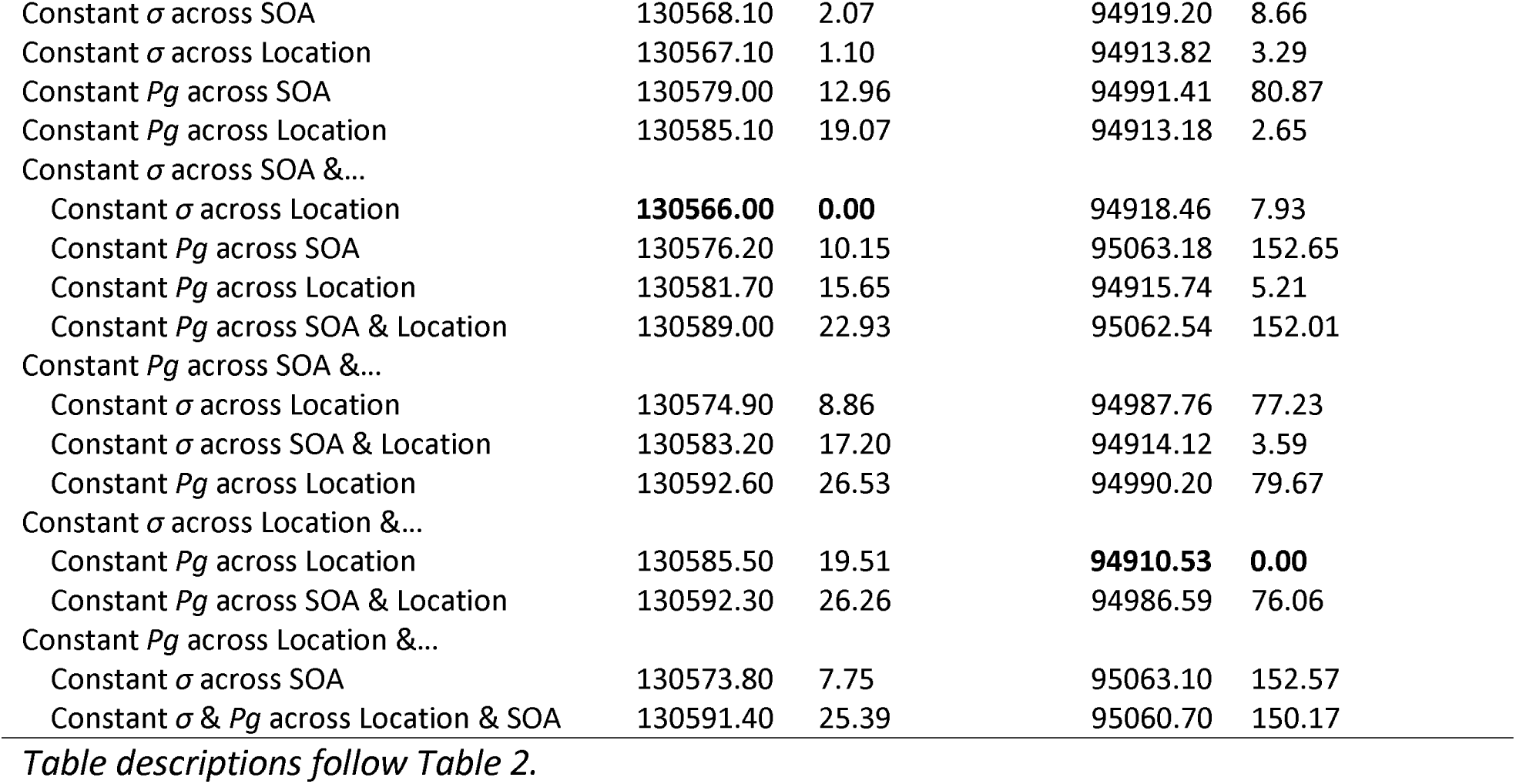
WAIC for all tested models in Experiment 4.

| Model | T1 |  | T2\|T1 |  |
| --- | --- | --- | --- | --- |
|  | WAIC | Difference from the best model | WAIC | Difference from the best model |
| Full model | 130569.60 | 3.58 | 94915.53 | 5.00 |
| Constant $\sigma$ across SOA | 130568.10 | 2.07 | 94919.20 | 8.66 |
| Constant $\sigma$ across Location | 130567.10 | 1.10 | 94913.82 | 3.29 |
| Constant $Pg$ across SOA | 130579.00 | 12.96 | 94991.41 | 80.87 |
| Constant $Pg$ across Location | 130585.10 | 19.07 | 94913.18 | 2.65 |
| Constant $\sigma$ across SOA &... | | | | |
| Constant $\sigma$ across Location | **130566.00** | **0.00** | 94918.46 | 7.93 |
| Constant $Pg$ across SOA | 130576.20 | 10.15 | 95063.18 | 152.65 |
| Constant $Pg$ across Location | 130581.70 | 15.65 | 94915.74 | 5.21 |
| Constant $Pg$ across SOA & Location | 130589.00 | 22.93 | 95062.54 | 152.01 |
| Constant $Pg$ across SOA &... | | | | |
| Constant $\sigma$ across Location | 130574.90 | 8.86 | 94987.76 | 77.23 |
| Constant $\sigma$ across SOA & Location | 130583.20 | 17.20 | 94914.12 | 3.59 |
| Constant $Pg$ across Location | 130592.60 | 26.53 | 94990.20 | 79.67 |
| Constant $\sigma$ across Location &... | | | | |
| Constant $Pg$ across Location | 130585.50 | 19.51 | **94910.53** | **0.00** |
| Constant $Pg$ across SOA & Location | 130592.30 | 26.26 | 94986.59 | 76.06 |
| Constant $Pg$ across Location &... | | | | |
| Constant $\sigma$ across SOA | 130573.80 | 7.75 | 95063.10 | 152.57 |
| Constant $\sigma$ & $Pg$ across Location & SOA | 130591.40 | 25.39 | 95060.70 | 150.17 |

#### Hypothesis Testing

Substantial and decisive evidence in favor of the main effect of SOA and Location was observed on T1 Pg (BF_10_ = 4.8; BF_10_= 186.8, respectively). T1 Pg averaged 0.27 in the short SOA condition and decreased to 0.24 in the long SOA condition (Table 1). Furthermore, T1 Pg averaged 0.28 in the same location condition and decreased to 0.24 in the different location condition. We found evidence against an interaction of SOA and Location on T1 Pg (BF_10_ < .1). Decisive and substantial evidence was observed for the main effect of SOA on T2|T1 Pg and σ (BF_10_> 1000; BF_10_ = 6, respectively). T2|T1 Pg averaged 0.46 at short SOA and decreased to 0.3 at long SOA (Figure 8c). Moreover, T2|T1 σ averaged 15.9° at short SOA and 14.9° at long SOA (Figure 8d).

### Discussion

The results were in line with the expectation that the introduction of a spatial factor in the classic RSVP task should elicit a degree of gradual awareness during the AB, as both T2|T1 Pg and σ were currently influenced by SOA. The finding of a precision effect in the current task also demonstrated that having a distractor stream does not by itself produce discrete awareness. Somewhat counterintuitively, the analyses furthermore showed that report precision in the condition in which target locations were not the same was not different from the condition in which they were. This might indicate that it is rather the overall allocation of attention across multiple locations, or the extent of attention, that may induce gradual awareness, rather than the actual switch of attention from one location to the other.

## Experiment 5

The collective evidence from the experiments so far has implicated space as a factor in gradual awareness during the AB. Precision effects were observed in Experiments 1, 2, and 4, all of which had targets appear across multiple locations. There was limited evidence that increasing the number of possible target locations, beyond more than a single one, increased the precision effect (cf. Experiment 1A and 1B). Conversely, the actual switch from one target location to another did not seem to modulate precision (Experiment 4). It may thus be hypothesized that the extent or distribution of attention, rather than its locus, is critical to the processing of T2, and determines whether its representation is consequently fully or only partially lost. This means that if the extent of the attentional ‘zoom lens’ is broad (i.e., covering more than a single location), gradual awareness will occur, regardless of whether the locus of attention moves across the visual field or not. However, this should also mean that if the extent of attention is narrow (i.e., covering a single location), then discrete awareness should occur, again regardless of whether attention moves.

To put this idea to the test, we implemented an RSVP design in which the extent of the attentional zoom lens that was induced by T1 was limited to a single location. To this end, each trial started with a single stream, containing T1 and a trailing distractor. Only then two lateral streams commenced, one of which contained T2. If our hypothesis is correct, then the narrow extent of the attentional zoom lens should produce a guess rate effect, that is, discrete awareness, even though attention moves on every trial to find T2.

### Method

#### Participants

Twenty-five students participated in the study in exchange for course credits or monetary reward. Two participants were excluded from analyses due to chance level error, so that 23 remained (15 females, 8 males; mean age = 22.7, range = 19-29). Procedure and design

Experiment 5 was identical to Experiment 3 with the following differences. The trial sequence was identical to that of Experiment 3, until the T1+1 distractor item (Figure 6c). After the T1+1 distractor, the single stream was split into two streams. T2 appeared either on the left or right stream, which was random, but evenly distributed in each condition. The spatial locations of the dual streams holding T2 were identical to those of Experiment 4. Depending on the condition, there were a total of 2 or 7 distractors between T1 and T2. There were a total of 408 trials per subject, so that there were 204 trials per SOA (lag) condition. Results Error: Bayesian paired-sample T-tests results showed anecdotal evidence against the effect of SOA on T1 error (BF_10_= 0.77; Table 1). In contrast, very strong evidence in favor of the effect of SOA on T2|T1 error was observed (BF_10_= 57.7). T2|T1 error (Figure 9a, b) averaged 29.8° at short SOA and decreased to 25.4° at long SOA. Model Comparisons: Model comparisons revealed that constant Pg across SOA was the best model explaining T1 error (Table 8). It should be noted that all other models also explained T1 error relatively well (WAIC _difference_ < 1.5). On the other hand, the constant σ across SOA model was the best model for T2|T1 error, replicating the results of Experiment 3 (Figure 9c, d).

**Figure 9.**
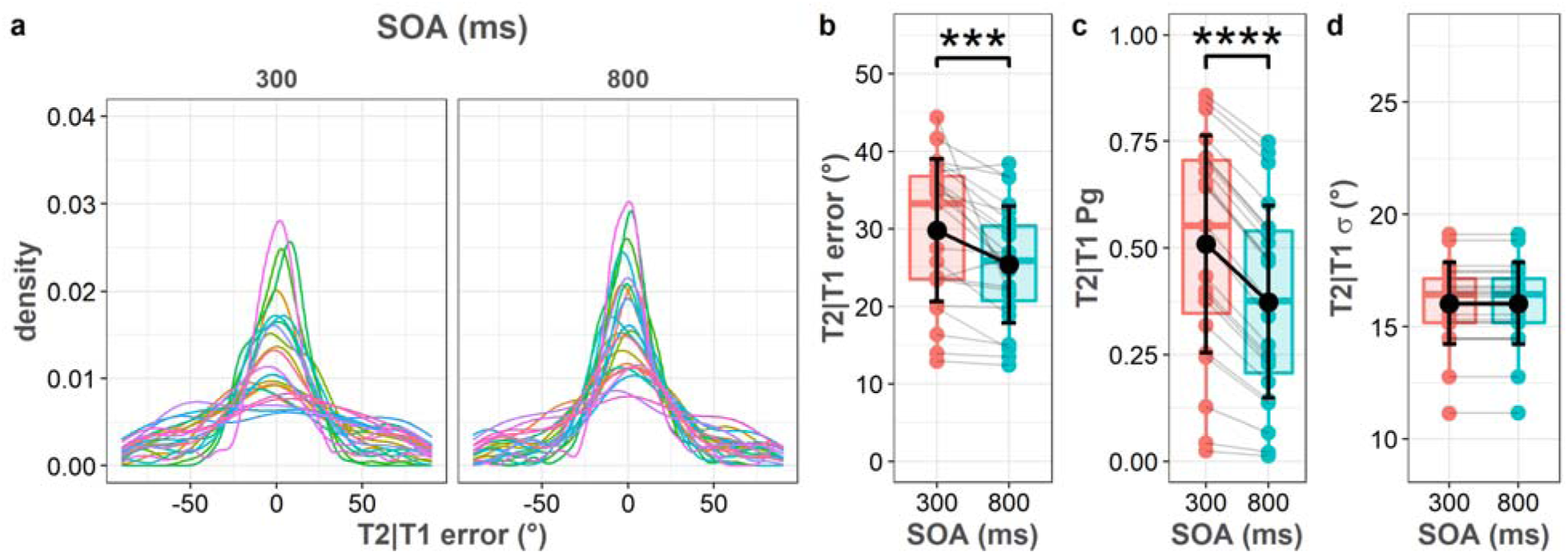
T2|T1 performance in Experiment 5. **a** Probability density plot of the T2|T1 error distribution of each subject for each SOA condition. **b** T2|T1 error, **c** T2|T1 Pg, **d** T2|T1 σ for short and long SOA conditions. Figure conventions follow Figure 2.

**Table 8.** WAIC for all tested models in Experiment 5.

| Model | WAIC | T1 | WAIC | T2\|T1 |
| --- | --- | --- | --- | --- |
|  |  | Difference From the |  | Difference From the |

|  |  | Best Model |  | Best Model |
| --- | --- | --- | --- | --- |
| Full Model | 99828.87 | 0.08 | 70452.07 | 1.69 |
| Constant $\sigma$ across SOA | 99829.67 | 0.89 | **70450.38** | **0.00** |
| Constant $Pg$ across SOA | **99828.78** | **0.00** | 70493.79 | 43.42 |
| Constant $\sigma$ and $Pg$ across SOA | 99830.09 | 1.31 | 70506.43 | 56.05 |

#### Hypothesis Testing

Bayesian tests revealed substantial evidence in favor of the effect of SOA on T1 σ (BF_10_ = 4.0). T1 σ averaged 15.7° at short SOA and decreased to 15.1° at long SOA. Decisive evidence was observed for the effect of SOA on T2|T1 Pg (BF_10_ > 1000). T2|T1 Pg averaged 0.51 at short SOA and decreased to 0.37 at long SOA (Figure 9c).

### Discussion

The outcomes were clear-cut: There was no evidence for gradual awareness in this task. There was evidence for a guess rate effect only, supporting the idea that awareness was discrete. Thus, it seemed that the narrow extent of the attentional zoom lens for T1 resulted in an all-or-none perception of T2, even though T2 did not appear within the same place as T1 or within its attentional boundaries. This outcome mirrors that of the classic single-stream RSVP in Experiment 3, and stood in contrast to the results of all other experiments, in which gradual target awareness was observed.

## General Discussion

As the title of the present article suggests, attentional selection can lead to two different faces of awareness; both gradual and discrete. In Experiments 1 and 2, which were classic DT tasks, the AB did not only modulate guess rates (a sign of discrete awareness), but also affected the representational precision of orientation and color stimuli. Experiment 3 showed that this was not true for the classic, single-stream RSVP task: The AB was expressed in elevated guess rates only. The hybrid RSVP designs implemented in Experiment 4 and 5 further elucidated the source of this difference in perceptual awareness. The dual-stream RSVP in Experiment 4 replicated the precision effect, as well as the guess rate effect, while the single-to-dual stream RSVP in Experiment 5 in turn replicated the guess rate effect only. Taken together, the results thus show cases of gradual and discrete awareness during the AB, with the dominance of either kind seemingly governed by the extent of the attentional zoom lens directed at T1. A narrow zoom lens, covering only a single location, triggers a specific attentional mechanism that ultimately causes all traces of blinked targets to perish from awareness. Conversely, a broad zoom lens, covering more than one location, allows a more variable degree of target-related information to enter awareness. Below, we discuss the implications of the current outcomes for existing theories of (temporal) attention and conscious awareness, followed by a new proposal for a conceptual model to characterize attentional gating of conscious awareness.

### Temporal attention

The current results reinforce the notion that some attentional deficit, an AB, occurs across time as well as space, and in this sense it remains a general limit on our cognitive abilities. The urgent question raised by the current study is why the nature of that limit is different depending on the number of possible T1 locations. Models of the AB do not currently provide an answer to this question as they typically assume that the attentional deficit is uniform across space (Olivers & Meeter, 2008; Taatgen et al., 2009; Wyble et al., 2009). Although proposals have been advanced to characterize spatiotemporal interactions, either by attributing these to different sources of interference (Wyble & Swan, 2015), or by introducing a degree of lateral inhibition to the general dynamic (Kristjansson & Nakayama, 2002), these still leave the general nature of the AB unchanged. The current results strongly suggest that it is time to incorporate the spatial dimension in models of temporal attention and the AB; not just as a dimension across which blink strength might vary, but also one in which adaptive interactions can occur.

The currently observed spatial interaction with target awareness during the AB does interface with two processes that are already postulated in models of the AB: Exerting cognitive control (Taatgen et al., 2009), and creating distinct episodic events (Wyble et al., 2009). Starting with the first, the nature of target awareness became more gradual when T1 appeared on more than one location, regardless of whether the location of T2 was variable or not (Experiment 5), and regardless of whether T2 actually appeared on the same location as T1 (Experiment 4). This suggests a certain strategic element in the way in which attention is controlled, even though it seems fully implicit. To process T1 optimally, a narrow attentional focus is chosen that also affects further processing during the AB. This in turn relates to the second process (creating distinct events), because this narrow focus not only facilitates the spatial search for T1 by restricting its extent, but may also foster a protective attentional gating mechanism that prevents subsequent events (i.e., T2) from interfering with the processing of T1. The existence of such a protective inhibitory gate is predicted explicitly by the eSTST model of the AB (Wyble et al., 2009), where it serves to maintain the episodic distinctiveness of the targets. This is an appealing functional justification that is highly compatible with the current results. Interestingly, models of the AB also assume that a higher investment in T1 increases the inhibitory gating response (Olivers & Meeter, 2008; Wyble et al., 2009, Taatgen et al., 2009; Wierda et al, 2012), which would be expected when the system is focused more narrowly on the location of T1. The gating mechanism in these models nevertheless still needs to be expanded to cover also cases in which attention is spread more broadly, which apparently diminishes the gating strength, and which produces a more gradual target awareness. Doing so will likely also necessitate the ability to represent continuous target dimensions, since the models were originally built to work with discrete stimuli.

### Working memory

Another issue of interest in awareness during the AB is its possible relationship with working memory. The current results are at least compatible with the idea that this relationship exists. In particular, the fate of targets that suffer from a gradual rather than a discrete AB provide a clue. These targets do not survive unscathed, as the quality of their representations is markedly reduced, which indicates that there is a rate limitation that applies to the overall process. Thus, even if these T2s do manage to creep into the binding pool (or workspace, second stage, etc.), due to weaker attentional gating, the T2 representations still cannot be processed with full vigor, and become or remain imperfect. This fits with the idea that the AB is due to target competition in visual working memory, as previously proposed by Shapiro, Raymond, and Arnell (1994). Such competitive interactions are likely to be related to the process of consolidation (Jolicœur & Dell’Acqua, 1998), implicating that this memory consolidation process need not necessarily operate in an all-or-none fashion either. Whether this is truly the case is uncertain. Ricker and Hardman (2017) showed evidence for discrete consolidation in visual working memory, and linked its time course directly to the AB, which fits with previous work from our group (Akyürek et al., 2007). However, even in the study of Ricker and Hardman (2017), there was at least “ambiguous evidence” for a gradual (precision) effect as well. Although the authors may have been right to view the discrete characteristic as the most important one in their design, it is conceivable that this may shift depending on task properties (e.g., stimulus timing, spatial layout, number of items to be recalled, etc.). Thus, more research is needed to further assess the relationship between working memory consolidation and the AB, and the nature of awareness therein.

### Spatial attention

Considering attentional processing more generally, one might speculate that the special status of location may reflect the primary spatial (retinotopic) organization of the visual cortex, particularly in earlier areas (Hubel & Wiesel, 1974). Electrophysiological studies have also consistently shown that space drives attention more directly than featural properties do (Anllo-Vento & Hillyard, 1996; Hillyard & Münte, 1984; Luck et al., 1994). It is therefore not surprising that influential models of attention have often assumed that space is a primary dimension, of a more fundamental nature than other features (such as color or orientation). For instance, in the Guided Search model (Wolfe, 1994), attention is driven by an activation map that places saliency signals coming from different feature dimensions on spatial coordinates. Similar spatial maps are implemented in other influential models of attention as well (e.g., Treisman &, Gelade, 1980; Itti & Koch, 2000; Bundesen et al., 2005). Despite the important role of space in these models, a temporally restricted interference dynamic, such as presently observed, is yet implicit in models of spatial attention, as they are typically concerned with single scenes, not successive events (though see also Petersen et al., 2012).

Another relevant aspect of spatial attention concerns the size of the attended area. A classic model of spatial attention, the ‘zoom lens’ model, correctly predicts that the size of the attended area is inversely related to perceptual performance (Eriksen & St. James, 1986). When limited attentional resources are spread out over a large attended area, they inadvertently lose their potency to facilitate stimulus processing. These effects are also evident from neuroimaging: When the size of the attentional field is large, the retinotopic area of preparatory neural activity increases, whereas the magnitude of this preparatory activity is attenuated (Müller et al., 2003). Here we found that the number of possible locations for T1 has clear effects on whether discrete and/or gradual awareness occurs during the AB, suggesting a possible relationship with the spatial trade-off predicted by the zoom lens model.

### Conscious awareness

The current finding that the AB can be expressed both as a discrete and as a gradual phenomenon has clear implications for theories of conscious awareness that put discrete selection during the AB at the center of that state of mind, such as the Global Neuronal Workspace model (Asplund et al, 2014; Dehaene et al., 2003; Sergent & Dehaene, 2004; Sergent et al., 2005). Clearly, a discrete AB is not necessarily typical for the nature of awareness, if observers are able to report at least partial information about blinked targets, as we primarily observed in our DT experiments. Rather, the discrete nature of awareness seems more due to a location-specific shielding mechanism that is presumably attentional in nature. It seems strenuous to suppose that awareness should be similarly location-specific, or that different global workspaces may be instantiated for different locations. In other words, a location-specific mechanism is hard to reconcile with a fundamental property of awareness, even though it is not out of place as a characteristic of attentional selection.

It must be stressed that the current results do not falsify the concept of a global workspace itself. The idea that an item in the center of awareness propagates through a large functional network, by which it claims a certain exclusivity at the expense of other pending items, is if anything strengthened by the currently observed blinks in both DT and RSVP tasks. It is rather the inhibitory gating mechanism, the binary choice between perceiving something and missing something completely, that seems to be less universal than previously believed, and which seems to depend on the extent of the attentional focus induced by T1. It may be hypothesized that even though conscious awareness may well be instantiated through a global neuronal workspace, this state of mind is distinct from the attentional mechanisms that deliver its contents. Thus, although attention may be binary in some cases and deliver either ‘perfect’ representations, or no representations at all, in other cases it may also provide a more steady stream of gradually impoverished ones.

More generally, it may be noted that literature about the nature of conscious awareness tends to classify awareness as if it has to be either discrete (e.g., Asplund et al., 2014; Tang et al., 2020; Dehaene et al., 2003; Sergent et al., 2005), or gradual (e.g., Overgaard et al., 2006; Ramsøy & Overgaard, 2004; Sandberg et al., 2010). The current outcomes showed that awareness does not necessarily have to be either one. Instead, we observed that within a single task, the nature of awareness can be both gradual and discrete, depending on momentary fluctuations. To better understand conscious awareness, research in this field might therefore rather focus on experimental manipulations that push the nature of awareness in either direction, instead of trying to assess what the nature of awareness is in any given case.

### The Adaptive Gating Hypothesis

To integrate the findings of the present study into an overarching framework in which both attention and conscious awareness are integrated, we developed a conceptual model that we labelled the “Adaptive Gating Hypothesis”. To recap, the current outcomes showed that the appearance of T1 not only triggers the well-known AB, but also determines the manner in which T2 enters conscious awareness. Awareness during the AB became more gradual when the observers had to monitor more than a single location or item (we use these terms interchangeably here) to find T1, which required them to increase the extent of their attentional zoom lens. The Adaptive Gating Hypothesis proposes that this change in the nature of awareness occurs because of an adaptive response to the resultant distribution of attentional resources that is needed when the location of T1 is variable.

As shown in Figure 10a, when attention is divided across more than a single item or location, there are fewer attentional resources available for each of them. These resources may reflect neural properties such as activity level (Müller et al., 2003), sampling frequency (e.g., VanRullen, 2016), or degree of pre-potentiation of, or recurrency within, the neural networks that are involved (Lamme & Roelfsema, 2000; Mashour et al., 2020). Regardless of the specific neural mechanism, the consequence is that target processing is impaired, compared to a situation in which T1 always appears at a single location. Indeed, there was some indication of comparatively higher error rates already on T1 itself when its location was variable (cf. Experiment 3 and 4). When the cognitive system is faced with a reduced ability to process the targets, it would be an adaptive response to tailor its processing mode to best meet these circumstances. We suggest that this is done by adjusting (i.e., reducing) the strength of the attentional gate, which is effectively a threshold that representations must pass to reach awareness (cf. Dehaene et al., 2006; van Vugt et al., 2018). Thus, when target processing is more difficult, due to spatial resource sharing, a lower threshold is set to help salvage as much target-related information as is available.

**Figure 10.**
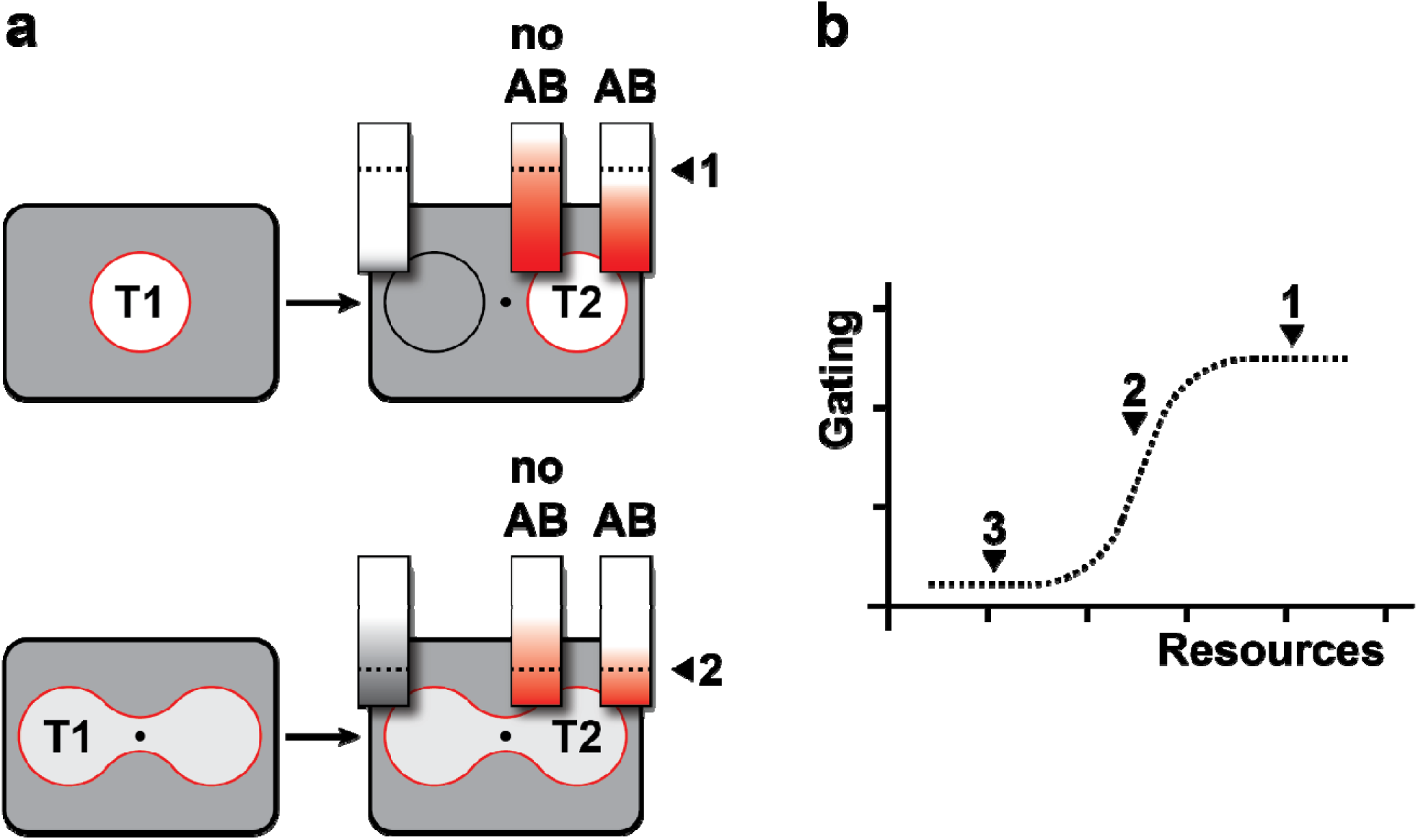
The Adaptive Gating Hypothesis. **a** Attention across target locations induced by thee spatial configuration of T1. Highlighted areas outlined in red show attentional allocation to aa single location (top) and multiple locations (bottom). The shape of the areas is arbitrary and for illustration purposes only. Gradient inset bars depict the amount of resources available at eachh location. For T2, the dual red gradient bars show resource levels in trials with and without an ABB (on the right and left, respectively). Dashed horizontal lines within indicate the hypothesized strength of the attentional gate; higher for a single location (top) and lower for multiple locations (bottom). **b** Attentional gating as a function of the amount of resources availablee. Point 1 indicates high gating strength when attention is allocated to a single location. Point 22 indicates low gating strength when attention is allocated to multiple locations. Point 3 indicatess absence of attentional gating when (almost) no attentional resources are needed (hypothetical single-target scenario).

As shown in Figure 10b, we hypothesize that gating strength follows a typical psychometric response function. This simple function not only characterizes a range of processes in both behavior and brain, from color discrimination responses to the firing rate of neurons (e.g., Hodgkin & Huxley, 1952), but has also been observed for object recognition in masking paradigms with short stimulus durations (Grill-Spector et al., 2000), and as such seems parsimonious as well as fitting. To account for the change in access to awareness that we observed in our experiments, we assume that gating strength is a function of the attentional resources that are allocated at any one location. We assume that the spatial properties of T1 initially modulate gating strength, but the associated effects are observed primarily on T2 during the AB, because it is then impacted by the processing of T1, and paying attention to T2 is consequently most difficult.

In more detail, according to our model, when T1 can only occur at a single location, the extent of the attentional zoom lens is small and resources can be concentrated to a small region in the visual field. Accordingly, gating strength is high (labeled 1 in the figure). Strong gating will assure that only well-processed targets (T2s) are selected and able to enter awareness, prioritizing selectivity and the quality of the representation, and producing an all-or-none response. Conversely, when T1 can occur at several locations, attention is spread across more than one location, and fewer resources are allocated at each of these locations. In this case, gating strength drops rapidly (labeled 2 in the figure). As indicated, this provides an important benefit: Since it is unlikely that under these circumstances a ‘perfect’ target representation will arise, a weaker gate ensures that at least some information is passed on to awareness. This prioritizes the quantity of information when it is scarce, but thereby also generates a more gradual target awareness. Lastly, in more extreme cases, that is, when very few attentional resources are allocated, gating strength is almost zero (labeled 3 in the figure). This scenario was not explicitly tested in the current AB tasks, but should apply in cases where no selection has to be made, such as in (masked) single-item paradigms, in which conscious awareness appears to be highly gradual indeed (Bar et al., 2001; Overgaard et al., 2006; Ramsøy & Overgaard, 2004; Sandberg et al., 2010).

### Open questions

The Adaptive Gating Hypothesis generates several testable predictions that might be further investigated to put this model to the test. First, our data suggest that the spatial properties of T1 induce the attentional configuration that applies to T2 as well (see also Jefferies et al., 2007 for related evidence in the context of Lag 1 sparing). When the extent of the attentional zoom lens induced by T1 is large, it seems likely that it will cover more than only the locus of T2. Given that in this case gating strength is low, other reportable items appearing in the vicinity of T2, such as distractors, may also break into awareness. The adaptive gating hypothesis also predicts the opposite pattern for cases in which attention is narrowly deployed; there should be almost no measurable awareness of items other than T2. Second, since the adaptive gating hypothesis holds that reduced attentional resources are compensated for by weaker gating, it follows that other manipulations that put a burden on attentional processing (as in Olivers & Nieuwenhuis, 2006, Arend et al., 2006; Taatgen et al., 2009) may have similar effects. In other words, increasing attentional load should result in effects on recall precision more than on guess rate. Third, although we observed a relatively rapid drop in gating strength as soon as more than a single item or location was monitored that would suit the proposed psychometric function, the shape of this curve should be substantiated, for instance by manipulating the extent of the attentional zoom lens in finer spatial detail. Fourth, since awareness is presumably modality-unspecific, it would seem worthwhile to investigate the gating function in other modalities, such as hearing. Auditory blinks certainly occur (e.g., Duncan et al., 1997), but space is not as primary to audition as it is to vision. In audition, time may be more important (Näätänen & Winkler, 1999). It thus remains to be seen whether and when attentional gating occurs in the former modality, or whether the AB might be predominantly gradual when target location does not play an important role.

### Conclusions

We found that conscious awareness during the AB can have two qualitatively different faces, such that blinked items may be completely or only gradually lost for report. This finding suggests that models of temporal attention, which have attributed the AB to a discrete, inhibitory gating mechanism, will need to account for the spatial specificity of this mechanism, and explain how a gradually impoverished target representation may emerge from the AB. Furthermore, theories of conscious awareness, which have assumed that the AB is exclusively discrete, and which have taken that as indicative of the nature of awareness, may thereby have been misguided. Finally, the Adaptive Gating Hypothesis provides a first account and a testable framework for the occurrence of both gradual and discrete awareness, but awaits further quantification and validation.

## Acknowledgements

AK and EGA were supported by an Open Research Area grant (464.18.114) from the Dutch Research Council (NWO), the Deutsche Forschungsgemeinschaft (DFG), and the Economic and Social Research Council (ESRC). JW was supported by an Abel Tasman Talent Grant from the Graduate School of Medical Sciences of the University of Groningen and Shenzhen University. We are thankful to our reviewers, in particular Timothy Ricker for his help with the model comparisons, and Roberto Dell’Acqua and Kimron Shapiro for their constructive comments on the interpretation of our results.

## Footnotes

1 It has been argued more generally that so-called low-level visual features (such as color) give rise to graded conscious awareness, while high-level features (such as word identity) do not (e.g., Windey & Cleeremans, 2015). The present study does not seek to arbitrate this hypothesis, but it is important to underscore that none of our conclusions rest on a difference at the feature level: We obtained evidence of both graded and discrete awareness with the same feature set.

