## Supplementary Material for "Two faces of perceptual awareness during the attentional blink: Gradual and discrete"

**1. Model recovery analysis**

To validate our methods, we conducted a model recovery analysis. This analysis shows whether our methods can accurately identify which model data was generated and whether the true parameters can be accurately and precisely estimated. We applied model recovery analysis for the main effect of SOA and Location (cf. Experiment 4). Data was generated from a mixture of a uniform and a von Mises distribution, assuming a base level of Pg = 0.2 and K = 5 (σ = 13.6). Effect sizes for each condition were *Pg* = 0.2 and *K* = 1, assuming no interaction between SOA and Location. We generated 100 trials per condition for 20 subjects. First, for the main effect of SOA on *Pg* and *σ*, data was aggregated across Location conditions yielding 200 trials for each SOA condition. Each generated dataset was then fitted to each of these four possible models. This generated a set of 4 (true models) x 4 (fitted models) = 16 model fits. The WAIC difference between the best fitting model and each model ($\Delta WAIC$) was computed. We estimated the bias and precision of the estimates for population parameters and effects by extracting the MCMC chains and subtracting the true population parameters and effects. For the SOA models, $\Delta WAIC$ for the true model was consistently low across models, suggesting that model recovery is relatively reliable (Figure S1a-c). Second, we fitted the same data for experimental designs with two main effects: SOA and Location. Data was generated for 16 models, varying the presence or absence of effects in *Pg* and *σ* in both SOA and Location conditions. Each generated dataset was then fitted to each of the 16 possible models. The model recovery analysis generated a set of 16 (true models) x 16 (fitted models) = 256 model fits. For the SOA X Location models, $\Delta WAIC$ for the true model was consistently low across models (2.13), again suggesting that model recovery is relatively reliable (Figure S1d-f). However, other $\Delta WAIC$ scores were often small too, favoring models that were more complex than the true model. This may result from common variance explained by *Pg* and *σ*, especially when *σ* is high (as *σ* tends to infinity, the von Mises distribution approximates a uniform distribution). Despite this bias, the effects of *Pg* and *σ* were not confused (i.e., there was a bias towards models where *both* parameters varied, not where only the *other* parameter varied; see middle four tiles in figure S1a).

Even though our methods were slightly biased towards more complex models, parameter recovery was relatively accurate when the full model was used. The mean and standard deviation of the errors in population parameters *Pg* (Figure S1b, S1e) and *σ* (Figure S1c, S1f) across simulated models was low (with a consistent underestimation of population parameter *σ*, possibly due to the default priors that were used). More importantly, differences between true and estimated effect sizes for *Pg* and *σ* across simulated models were also consistently low and within the range of the true effect sizes. In sum, this analysis suggests that our methods are able to recover both the true model and underlying parameters of simulated data relatively well, despite small biases to more complex models in the $\Delta WAIC$ scores. All model recovery results are available in the OSF repository that is referenced in the main text.

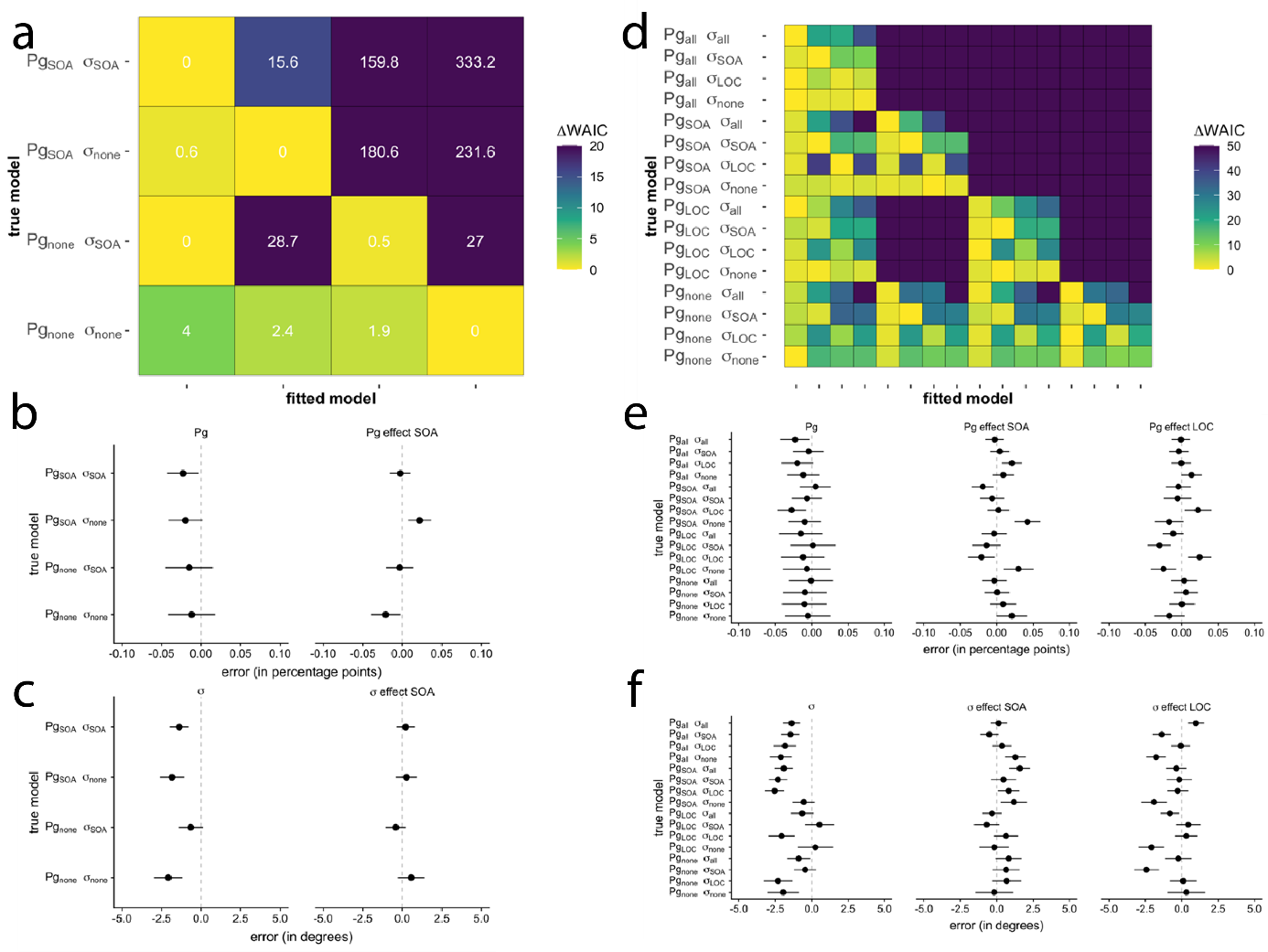
***Figure S1.*** *Model recovery analysis.* ***a*** *Model recovery analysis for models where only SOA was varied. Colors indicate the difference in WAIC between the best fitting model and the other models, with the true model on the x-axis, and the fitted model on the y-axis. Models are sorted according to complexity (top to bottom, left to right), illustrating that more complex fitted models (left) generally have lower* $\Delta WAIC$ *scores.* $\Delta WAIC$ *scores on the diagonal signify how well true models can be recovered.* ***b*** *Parameter recovery for Pg for models where only SOA was varied. For each true model (y-axis), the mean and standard deviation of the error (x-axis) are shown for both population parameters (Pg), as well as the effect size for SOA.* ***c*** *Parameter recovery for σ for models where only SOA was varied.* ***d*** *Model recovery analysis for models where both SOA and Location were varied.* ***e*** *Parameter recovery for Pg for SOA x Location models.* ***f*** *Parameter recovery for σ for SOA x Location models.*

**2. T1 Figures Across Experiments**

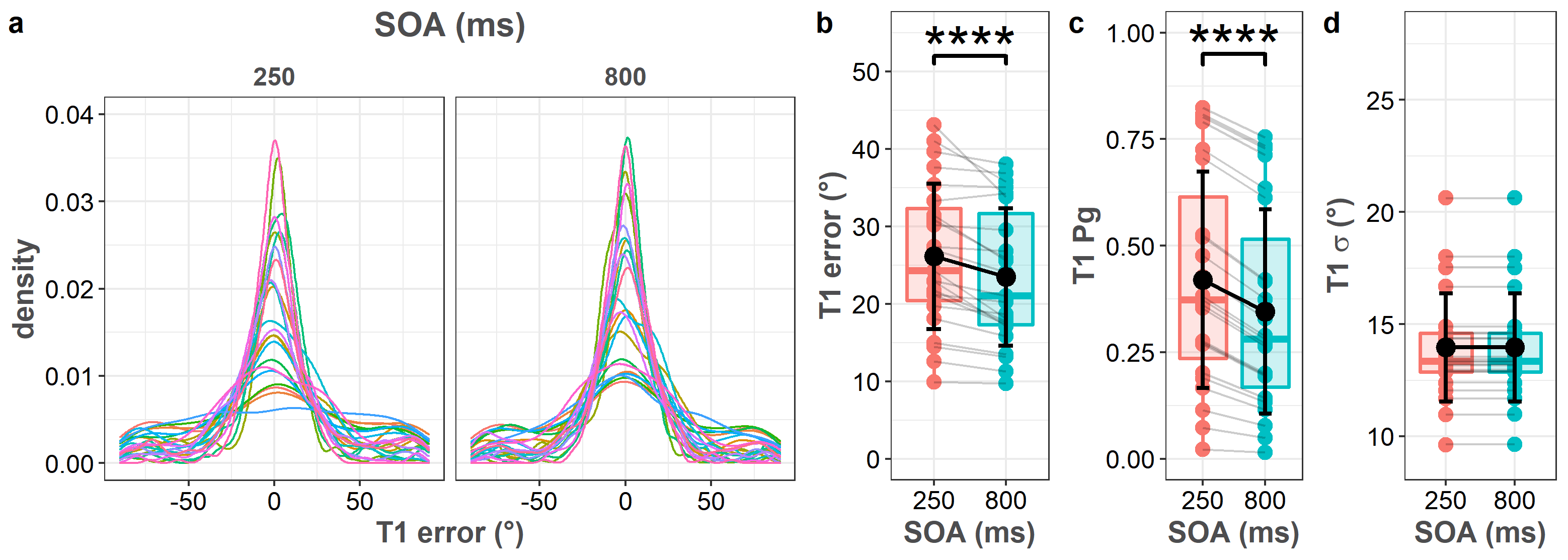
***2.1. Experiment 1A***

***Figure S2.*** *T1 performance in Experiment 1A.* ***a*** *Probability density plot of the T1 error distribution of each subject for each SOA condition.* *Each color indicates a subject.* ***b*** *T1 error for short and long SOA conditions (ms). Colored dots represent individual data, black dots reflect averages, and error bars represent 95% confidence intervals. Grey lines connect each subject in short and long SOA conditions. Whisker plots show median and quartile values. The orange color shows the short SOA condition and the blue color shows the long SOA condition. * indicates anecdotal evidence against the null hypothesis, ** indicates substantial evidence against the null hypothesis, *** indicates strong evidence, and **** means decisive evidence.* ***c*** *T1 Pg based on the best model predictions.* ***d*** *T1 σ based on the best model predictions.*

***2.2. Experiment 1B***

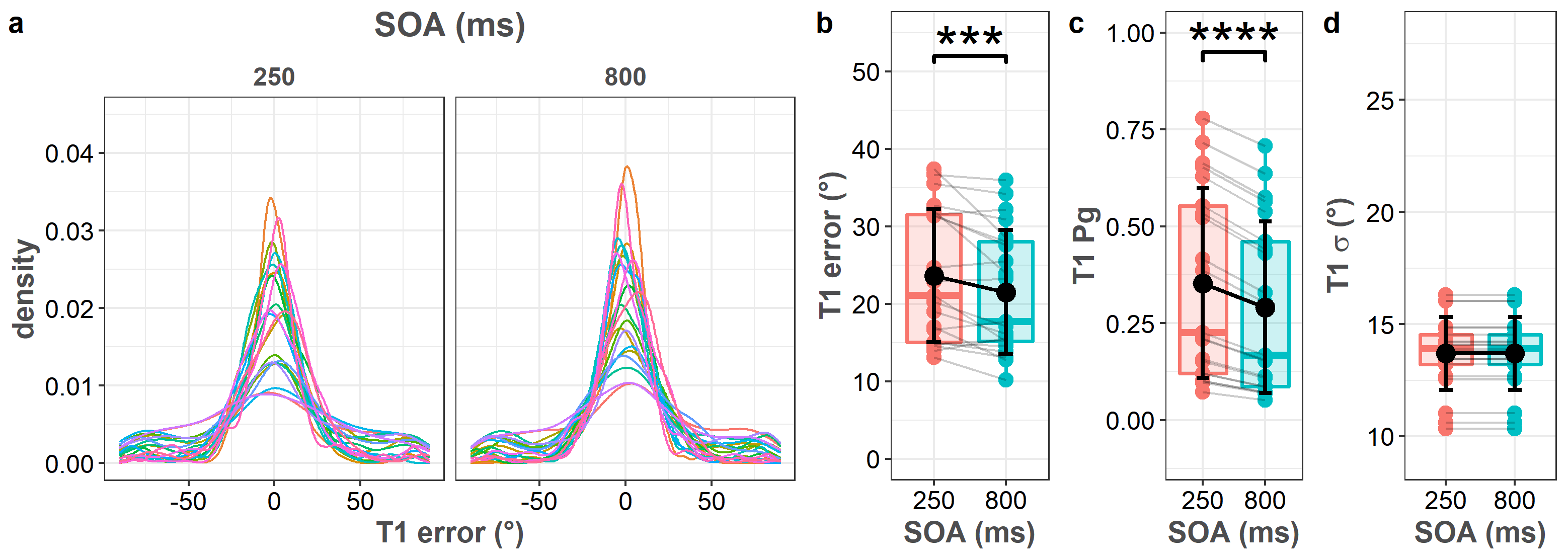
***Figure S3****. T1 performance in Experiment 1B.* ***a*** *Probability density plot of the T1 error distribution of each subject for each SOA condition.* ***b*** *T1 error,* ***c*** *T1 Pg,* ***d*** *T1 σ for short and long SOA conditions. Figure conventions follow Figure S2.*

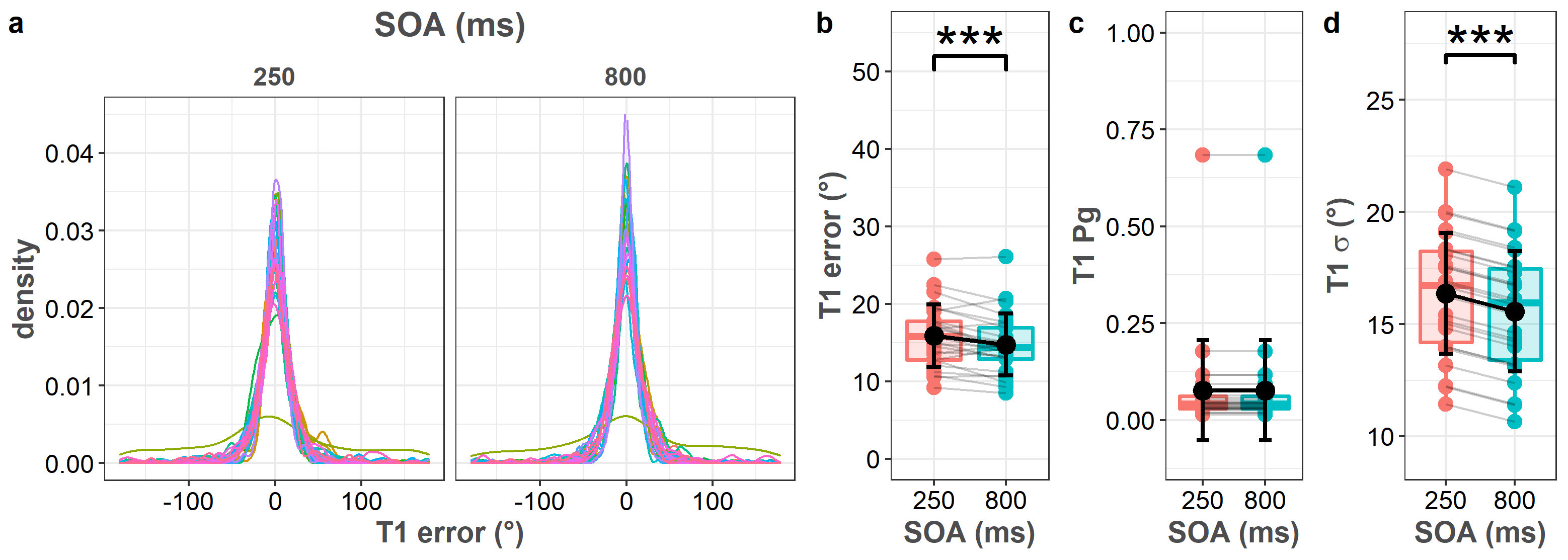
***2.3. Experiment 2A***

***Figure S4****. T1 performance in Experiment 2A.* ***a*** *Probability density plot of the T1 error distribution of each subject for each SOA condition.* ***b*** *T1 error,* ***c*** *T1 Pg,* ***d*** *T1 σ for short and long SOA conditions. Figure conventions follow Figure S2.*

***2.4. Experiment 2B***

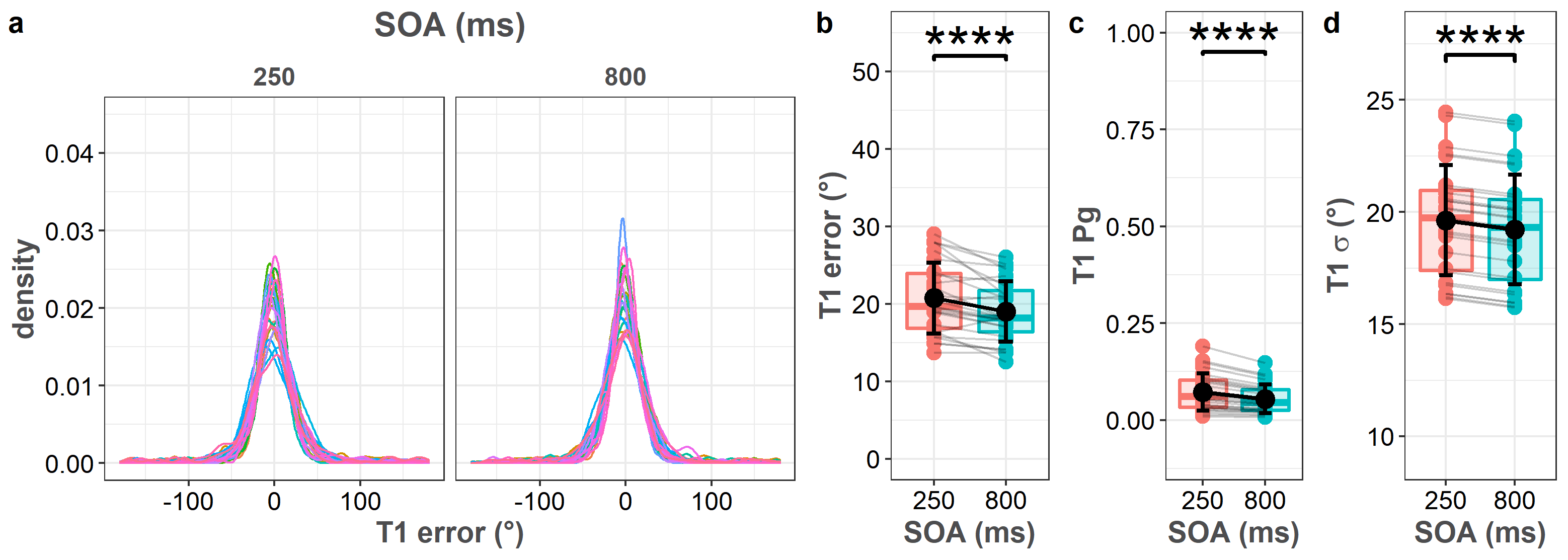
***Figure S5****. T1 performance in Experiment 2B.* ***a*** *Probability density plot of the T1 error distribution of each subject for each SOA condition.* ***b*** *T1 error,* ***c*** *T1 Pg,* ***d*** *T1 σ for short and long SOA conditions. Figure conventions follow Figure S2.*

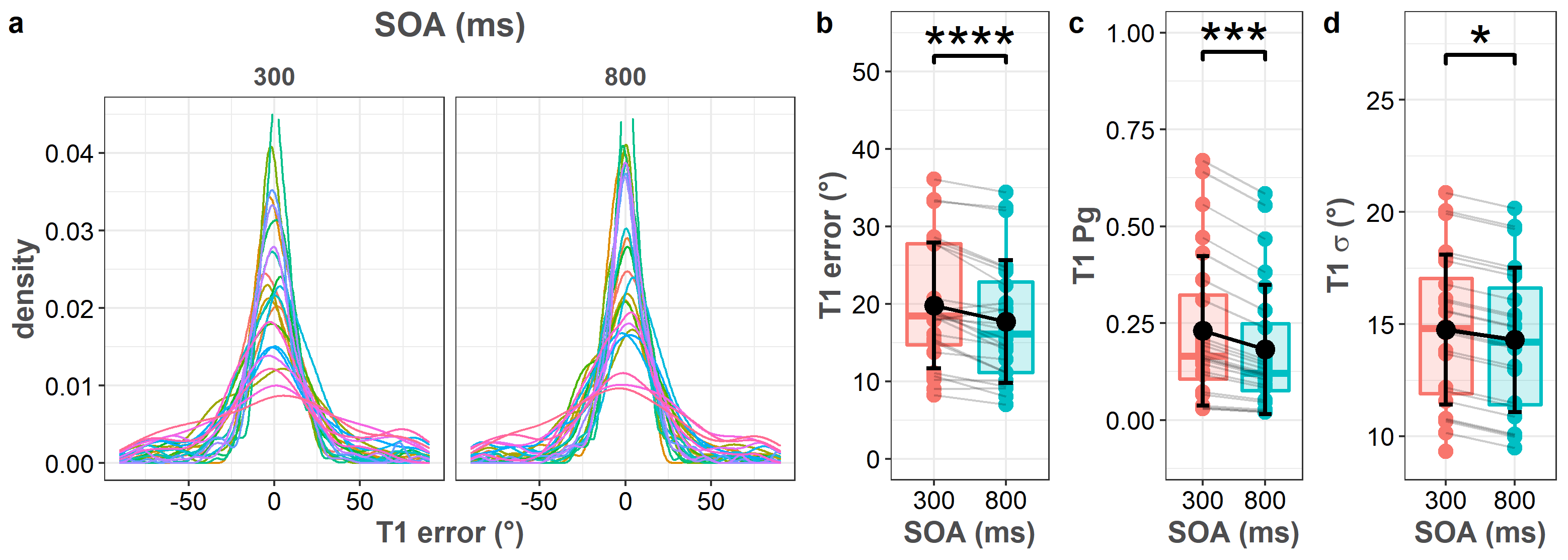
***2.5. Experiment 3***

***Figure S6****. T1 performance in Experiment 3.* ***a*** *Probability density plot of the T1 error distribution of each subject for each SOA condition.* ***b*** *T1 error,* ***c*** *T1 Pg,* ***d*** *T1 σ for short and long SOA conditions. Figure conventions follow Figure S2.*

***2.6. Experiment 4***

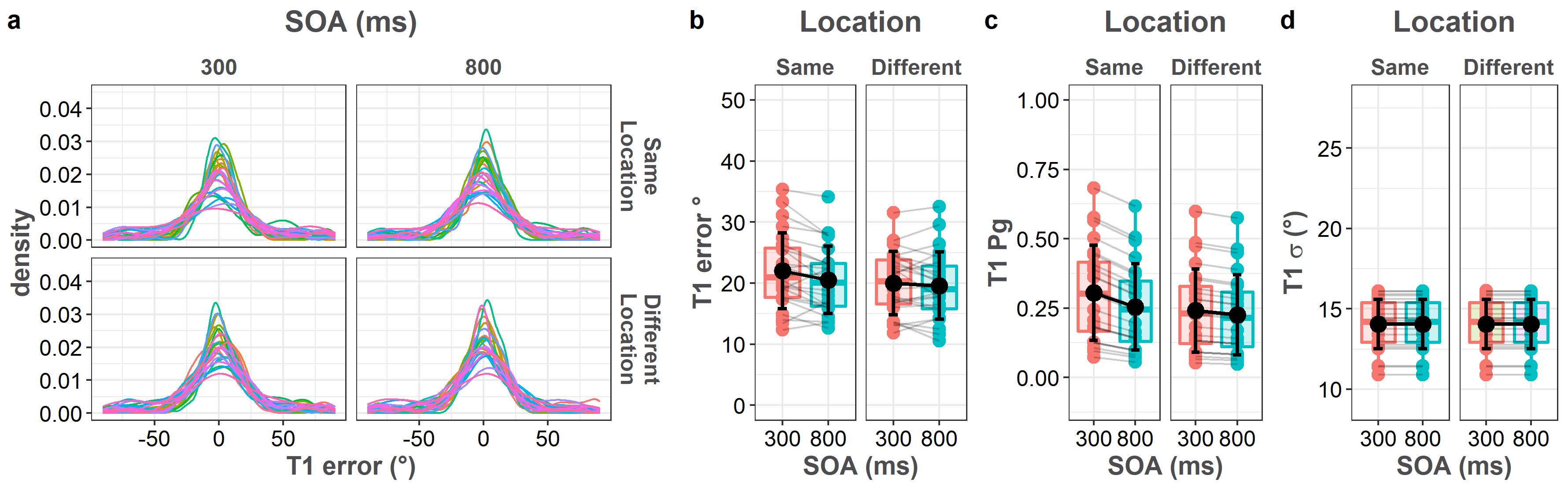
***Figure S7****. T1 performance in Experiment 4.* ***a*** *Probability density plot of the T1 error distribution of each subject for all SOA & Location conditions.* ***b*** *T1 error,* ***c*** *T1 Pg,* ***d*** *T1 σ across SOA & Location conditions. Figure conventions follow Figure S2.*

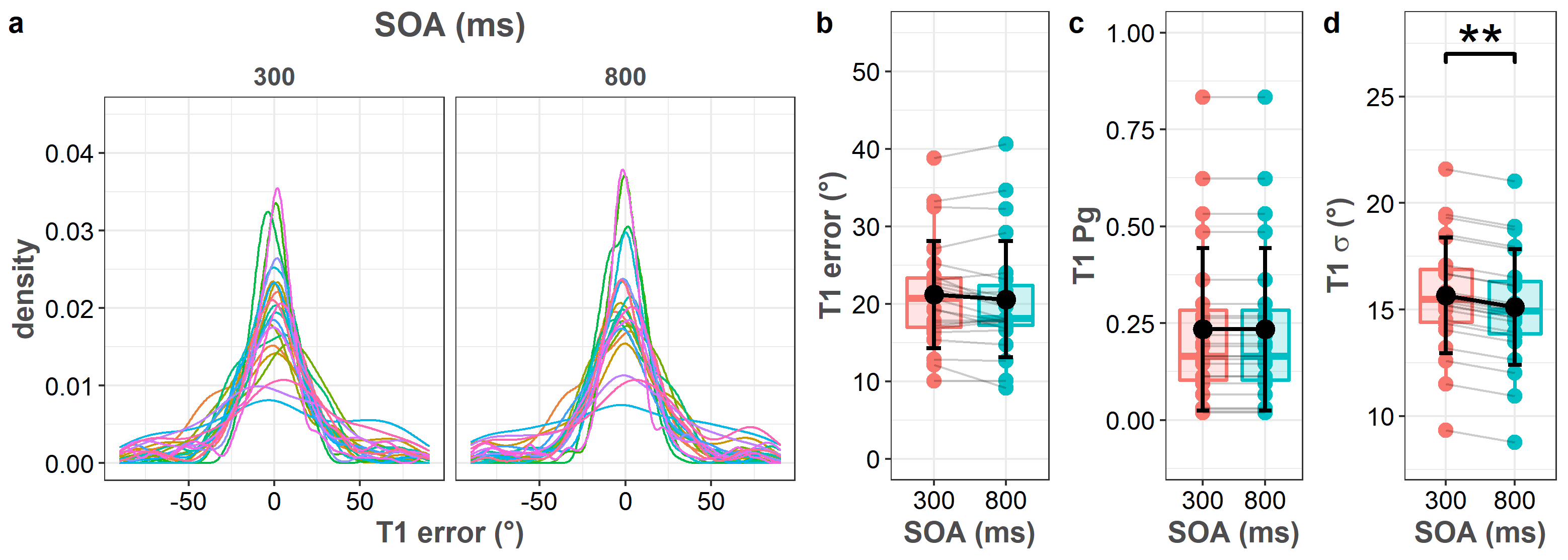
***2.7. Experiment 5***

***Figure S8****. T1 performance in Experiment 5.* ***a*** *Probability density plot of the T1 error distribution of each subject for each SOA condition.* ***b*** *T1 error,* ***c*** *T1 Pg,* ***d*** *T1 σ for short and long SOA conditions. Figure conventions follow Figure S2.*

**3. Experiment 4: Bayesian repeated-measures ANOVA results on T1 and T2|T1 error**

Bayesian ANOVA analysis revealed that the main effect of SOA explains both T1 and T2|T1 error better than other models (Table S1). For T1 error, the model including only the main effect of SOA was 1.6 times more likely than the model including both main effects of SOA and Location, 8.4 times more likely than the model with main effects of SOA and Location as well as their interaction, 675.3 times more likely than the model with only the main effect of Location, and 2026 times more likely than the null model. For T2|T1, the model including only the main effect of SOA was 15.5 times more likely than the model including both main effects of SOA and Location, 63.6 times more likely than the model with main effects of SOA and Location as well as their interaction, 8320987.6 times more likely than the model with only the main effect of Location, and 187482614.7 times more likely than the null model.

***Table S1****. Bayesian repeated measure ANOVA model comparison of the interaction of Location and SOA on T1 and T2|T1 error in Experiment 4.*

|  | **T1 error (°)** | | | | | **T2\|T1 error (°)** | | | | |
| --- | --- | --- | --- | --- | --- | --- | --- | --- | --- | --- |
| ***Models*** | ***P(M)*** | ***P(M\|data)*** | ***BF _M_*** | ***BF _10_*** | ***error %*** | ***P(M)*** | ***P(M\|data)*** | ***BF _M_*** | ***BF _10_*** | ***error %*** |
| Null model | 0.2 | 5.612e -4 | 0.002 | 1 |  | 0.2 | 1.798e -8 | 7.19e -8 | 1 |  |
| **SOA** | 0.2 | 0.503 | **4.052** | **896.729** | 1.606 | 0.2 | 0.771 | **13.48** | **4.290e +7** | 1.841 |
| LOCATION | 0.2 | 0.002 | 0.006 | 2.681 | 1.078 | 0.2 | 4.057e -9 | 1.62e -8 | 0.226 | 1.22 |
| SOA + LOCATION | 0.2 | 0.387 | 2.527 | 689.943 | 6.832 | 0.2 | 0.178 | 0.869 | 9.925e +6 | 3.093 |
| SOA ✻ LOCATION | 0.2 | 0.108 | 0.482 | 191.661 | 2.65 | 0.2 | 0.05 | 0.212 | 2.803e +6 | 2.431 |

*Note. All models include subject; SOA ✻ LOCATION = SOA + LOCATION + SOA:LOCATION; models are compared with the null model.*

**4. Bayesian hypothesis testing on full models of T2|T1 in Experiment 2A, 3, 4 and 5**

In Experiment 2A, 3, 4 and 5, restricted models produced the best fit, and therefore these were used in the main analyses for hypothesis testing. For reference, here we present hypothesis testing on guess rate and precision parameters based on the full models.

**4.1. Experiment 2A**

Strong evidence was observed against the main effect of SOA on T2|T1 *Pg (BF_10_* = .07, Figure S9a*).* T2|T1 *Pg* averaged 0.07 at short SOA and 0.07 at long SOA. Decisive evidence was observed in favor of an effect of SOA on T2|T1 *σ* (*BF_10_* > 1000, Figure S9b). Lastly, T2|T1 *σ* averaged 18.6° at short SOA and decreased to 17.2° at long SOA.

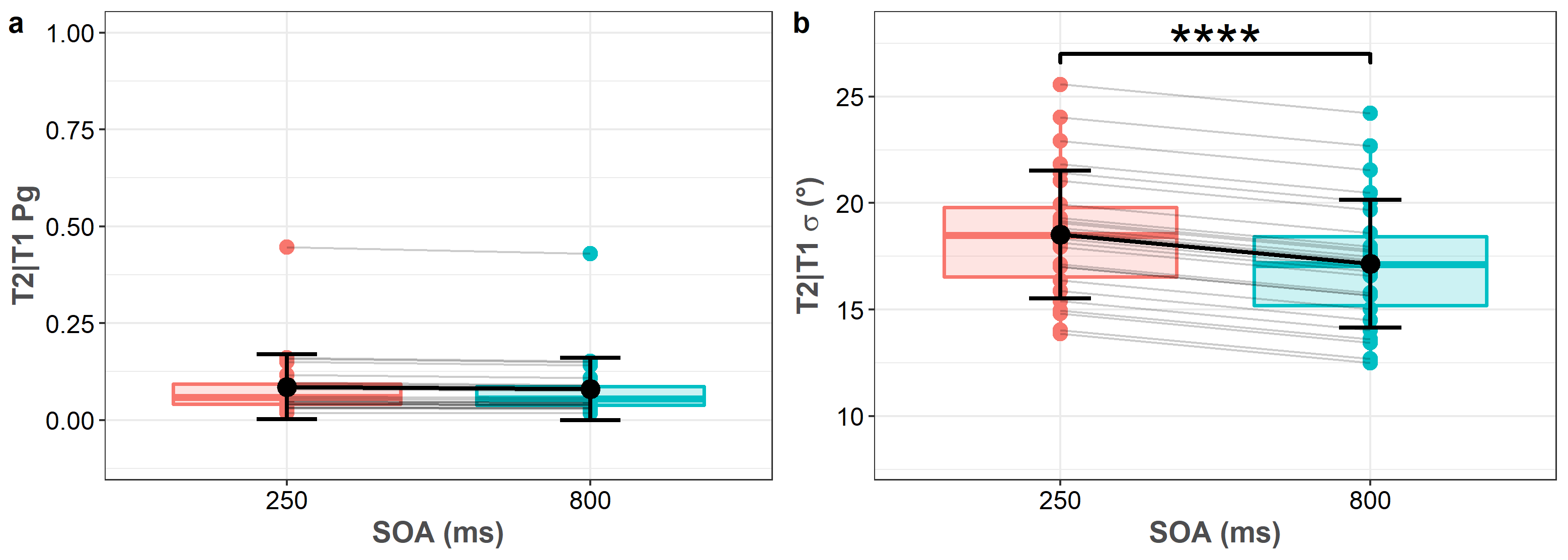

***Figure S9****. T2|T1 hypothesis testing with the full model in Experiment 2A.* ***a*** *T2|T1 Pg,* ***b*** *T2|T1 σ for short and long SOA conditions. Figure conventions follow Figure S2.*

**4.2. Experiment 3**

Decisive evidence was observed in favor of an effect of SOA on T2|T1 *Pg* (*BF_10_* > 1000, Figure S10a). T2|T1 *Pg* averaged 0.63 at short SOA and 0.12 at long SOA. Anecdotal evidence was observed against the main effect of SOA on T2|T1 *σ* (*BF_10_* = .62, Figure S10b). T2|T1 *σ* averaged 34.5° at short SOA and decreased to 34.0° at long SOA.

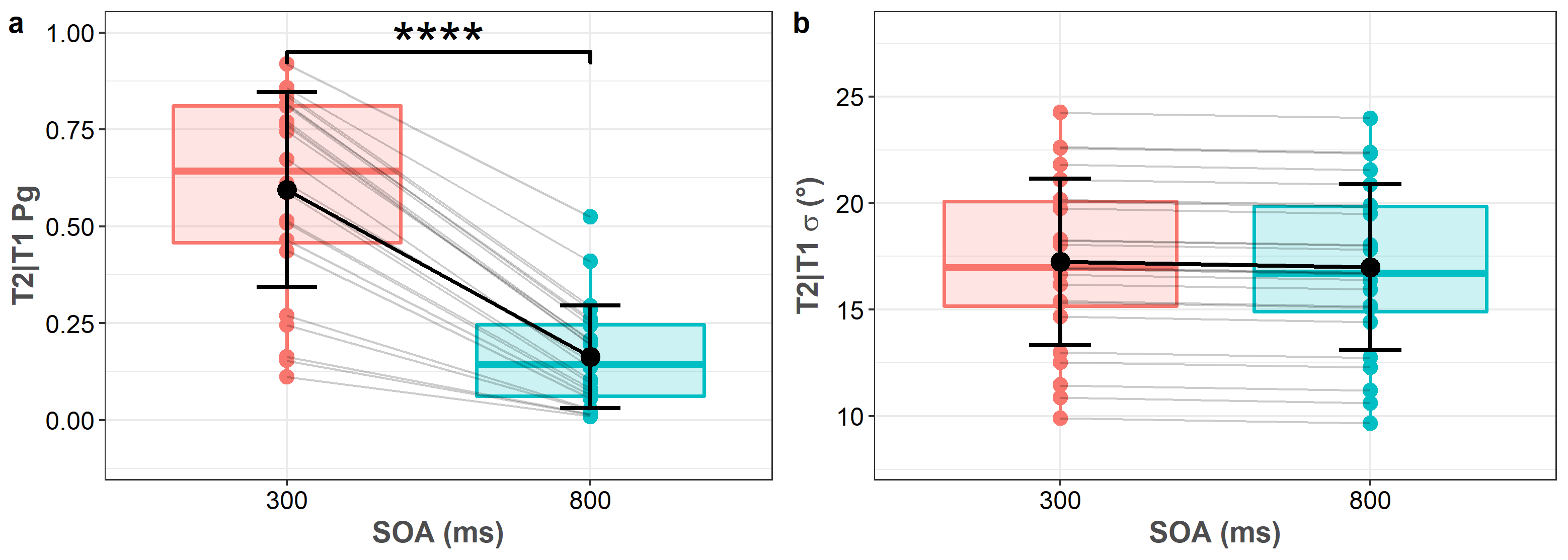

***Figure S10****. T2|T1 hypothesis testing with the full model in Experiment 3.* ***a*** *T2|T1 Pg,* ***b*** *T2|T1 σ for short and long SOA conditions. Figure conventions follow Figure S2.*

**4.3. Experiment 4**

Anecdotal and decisive evidence was observed for the main effect of SOA on T2|T1 σ and Pg (*BF_10_* = 2.3; *BF_10_* > 1000, respectively, Figure S11a-b). T2|T1 *Pg* averaged 0.46 at short SOA and decreased to 0.28 at long SOA. Moreover, T2|T1 *σ* was 15.9° at short SOA and 14.9° at long SOA. There was no evidence for the main effect of Location or for the interaction of Location and SOA on T2|T1 *Pg* or *σ* (*BF_10_* < 1).

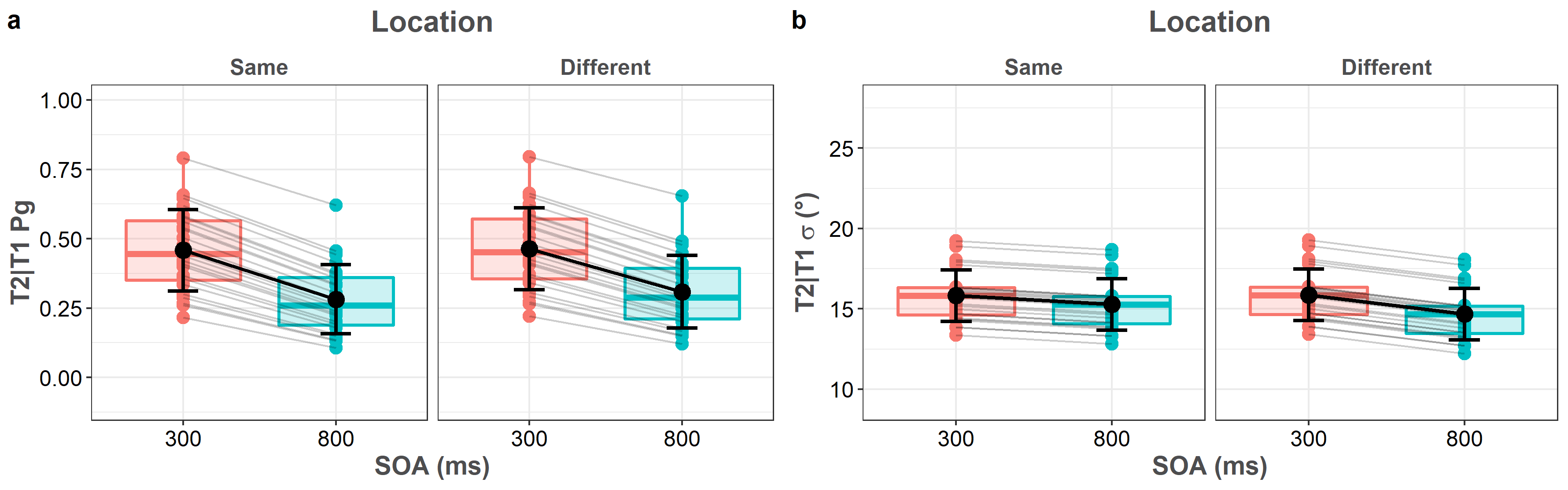

***Figure S11****. T2|T1 hypothesis testing with the full model in Experiment 4.* ***a*** *T2|T1 Pg,* ***b*** *T2|T1 σ for short and long SOA conditions. Figure conventions follow Figure S2.*

**4.4. Experiment 5**

Decisive evidence was observed in favor of an effect of SOA on T2|T1 *Pg* (*BF_10_* > 1000, Figure S12a). T2|T1 *Pg* averaged 0.49 at short SOA and decreased to 0.32 at long SOA. Anecdotal evidence was observed against the main effect of SOA on T2|T1 *σ* (*BF_10_* = .75, Figure S12b). T2|T1 *σ* was 16.1° at short SOA and 16° at long SOA.

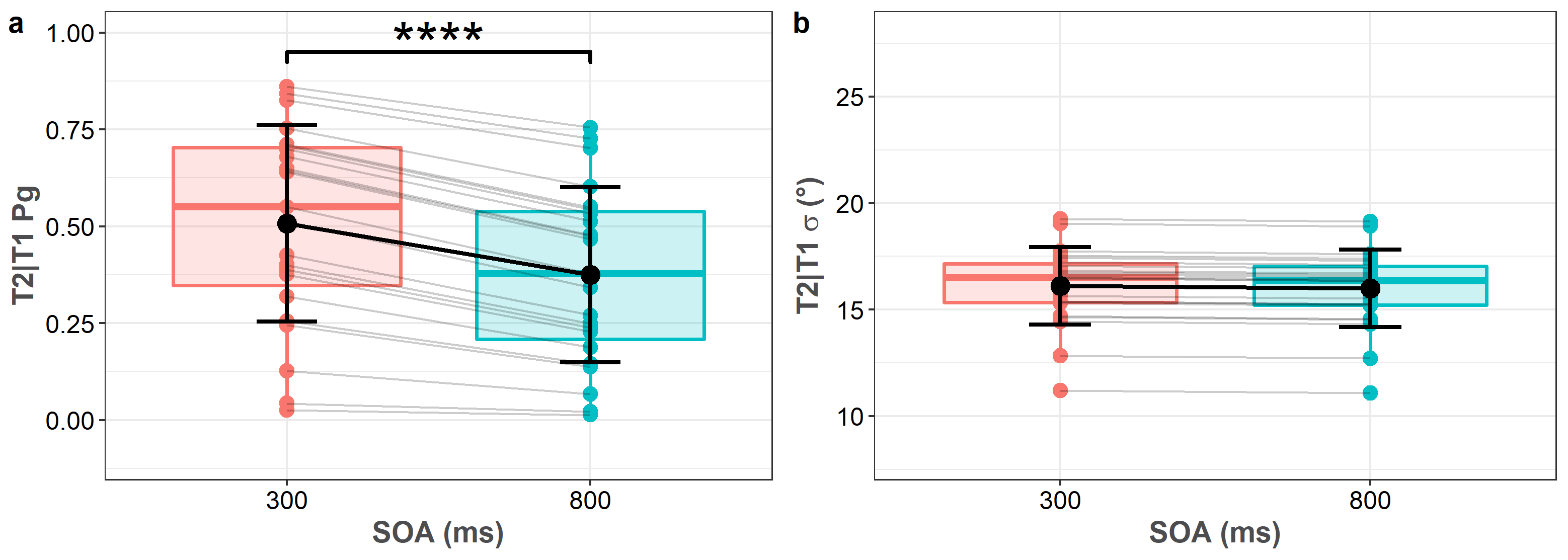

***Figure S12****. T2|T1 hypothesis testing with the full model in Experiment 4.* ***a*** *T2|T1 Pg,* ***b*** *T2|T1 σ for short and long SOA conditions. Figure conventions follow Figure S2.*
